# Development of non-β-Lactam covalent allosteric inhibitors targeting PBP2a in Methicillin-Resistant *Staphylococcus aureus*

**DOI:** 10.1101/2024.05.29.596450

**Authors:** Bogdan M. Benin, Rupak Kharel, Trae Hillyer, Chuanqi Sun, Anna Cmolik, Taylor Kuebler, Yuk Yin Sham, Robert Bonomo, Jeffrey D. Mighion, Woo Shik Shin

## Abstract

Methicillin-resistant *Staphylococcus aureus* (MRSA), a Gram-positive bacterial pathogen, continues to pose a serious threat to the current public health system in our society. The high level of resistance to β-lactam antibiotics in MRSA is attributed to the expression of penicillin-binding protein 2a (PBP2a), which catalyzes cell wall cross-linking. According to numerous research reports, the activity of the PBP2a protein is known to be regulated by an allosteric site distinct from the active site where cell wall cross-linking occurs. Here, we conducted a screening of 113 compounds containing a 1,3,4-oxadiazole core to design new covalent inhibitors targeting the allosteric site of PBP2a and establish their structural-activity relationship. The stereochemically selective synthesis of sulfonyl oxadiazole compounds identified in the initial screening resulted in a maximum eightfold enhancement in cell inhibition activity. The sulfonyl oxadiazole-based compounds formulated as PEG-based ointments, with low toxicity test results on human cells (CC_50_: >78μM), demonstrated potent antimicrobial effects not only in a mouse skin wound infection model but also against oxacillin-resistant clinical isolate MRSA (IC_50_ ≈ 1μM), as evidenced by the results. Furthermore, additional studies utilizing LC-MS/MS and in-silico approaches clearly support the allosteric site covalent binding mechanism through the nucleophilic aromatic substitution (S_N_Ar) reaction, as well as its association with the closure of the major active site of PBP2a.

## Introduction

Methicillin-resistant *Staphylococcus aureus* (MRSA) infections affect hundreds-of-thousands of Americans every year, resulting in additional $1.7 billion in attributable healthcare costs and tens-of-thousands of deaths ^1,2^. Although several new drugs have been approved in the past twenty years, there are few antibiotics currently in the clinical trial pipeline and mortality rates associated with MRSA infections have not significantly decreased ^3^. For this reason, researchers continue searching for new antibiotics based on structure types that differ from currently utilized drugs such as β-lactams, glycolipopeptides, etc.

All the *Staphylococcus aureus* strains have from PBPs (PBP1 to PBP4), but MRSA express a special PBP (PBP2 or PBP2a) from the mecA gene PBP2a takes over the biosynthetic function of normal PBPs in the presence of inhibitory concentration of β-lactams because PBP2 has a decreased binding affinity to β-lactams.

Recently, several groups have reported successful studies on the use of various heterocycle-based compounds, namely quinazolinones and oxadiazoles ^4-10^. The former were found to exhibit remarkably minimum inhibitory concentration (MIC) values as well as water solubility. Furthermore, they were demonstrated to inhibit penicillin binding protein 2a (PBP2a) allosterically. PBPs are involved in cell-wall synthesis and are generally the target of β-lactam antibiotics. MRSA, however, expresses PBP2a, which contains a closed active site until it has been allosterically activated by cell wall peptide fragments. This allosteric mechanism is targeted by the only two β-lactams with significant anti-MRSA activity: ceftaroline and ceftobiprole ^11^.

The same group that investigated the use of quinazolinones ^4,8,12^ also found that that oxadiazoles could also serve as antibacterial agents against MRSA ^6,7,13,14^. They found, through another screening and synthesis study, that the 1,2,4-oxadiazoles are also active against MRSAc^14^. The identified lead compound was 4-(3-(4-(4-(trifluoromethyl)phenoxy)phenyl)-1,2,4-oxadiazol-5-yl)aniline, which exhibited an MIC of 2 μg/mL (5 μM) ^13,14^. Although they were unable to confirm, crystallographically, that these compounds also bind to the same allosteric site as the quinazolinones or the 5th generation anti-MRSA cephalosporins ^15-17^, the authors could demonstrate potent synergism between β-lactams and this new class of antibiotics ^5^. This synergism is in all likelihood due to allosteric inhibition by the oxadiazole, followed by competitive inhibition by the β lactam ^5^. This work was followed by further research into this structure type ^18,19^, and several compounds with lower MIC values down to 0.5 μg/mL (1 μM) were reported ^20^. Although effective, some of these compounds exhibited noticeable cytotoxicity and their efficacy did not translate well to animal studies ^7,21^.

Importantly, the 1,2,4-oxadiazoles only represent one possible set of structural isomers, with 1,2,5-and 1,3,4-oxadiazoles also being investigated by medicinal chemists ^22^. Although both the 1,2,4-and 1,3,4-oxadiazoles are highly studied, scientists at AstraZeneca published a report indicating that the 1,3,4-isomers are expected to exhibit greater metabolic stability as compared to matched 1,2,4-oxadiazole regio-isomers ^22^. They suggest that this is in part due to decreased lipophilicity of 1,3,4 analogues as compared to their 1,2,4 counterparts, higher dipole moments, and decreased cytochrome P450 recognition ^22^. Since then, there have been only a few reports on the antibacterial properties of new 1,3,4-oxadiazoles ^23-26^. Many of these compounds are ineffective against MRSA, including those which were bound to fluoroquinolones. Of those which demonstrated potent antibacterial activity ^23,24^, these typically had an inert, aromatic group directly attached at C2 of the 1,3,4-oxadiazole ring and aryl-amide derivatives at C5 ^23^.

Based on these findings, this class remains open to further investigation as many other alternative substituent patterns have not been thoroughly considered. For example, the Shoichet group published a report detailing the identification of 2 sulfonyl-1,3,4-oxadiazoles as potential covalent β-lactamase inhibitors ^27^. Both compounds were found to be potent AmpC inhibitors with IC_50_ <1 μM, and the mechanism was suggested to be nucleophilic aromatic substitution with the sulfonyl substituent acting as the leaving group. Based on these promising results, and the lack of thorough follow up work, we set the 5-sulfonyl-1,3,4-oxadiazole ring and substituent as our template core from which we began our screening.

Herein, we report the computational screening, stereoselective synthesis and characterization of a new set of 5-sulfonyl-1,3,4-oxadiazoles, demonstrating their potent anti-MRSA activity in vitro and in vivo.

## Results and Discussion

Following several existing reports on oxadiazoles as non-β-lactam antibiotics with activity against various bacteria and drug-resistant enzymes, we utilized a commercially available compound library for the initial screening (**Supplementary Figure S1**). For this initial investigation, a list of 77 compounds were selected, including eight 5-thio-1,3,4-oxadizoles and sixty-nine 5-sulfonyl-1,3,4-oxadiazoles cores. The bacterial cell inhibition efficacy of these initial compounds (50 μM) was tested against MRSA (ATCC BAA-44, 1×10^8^ CFU/mL; **Supplementary Figure S1** and **Supplementary Table S1**) using a single-dose assay. Amoxicillin and meropenem were included as control groups, where MRSA exhibited significant resistance with survival rates of 85% and 81%, respectively, after 18 hours of treatment. Twenty-one of these compounds reduced the survival rate of MRSA cells by more than 80%, and the antibacterial activity of these 21 compounds was further characterized. Importantly, all 21 selected compounds have the same 5-sulfonyl-1,3,4-oxadiazole core structure, indicating that other heteroatoms severely inhibit activity To further understand the activity of the remaining 21 compounds, minimum inhibitory concentrations (MIC) were determined using the same ATCC BAA-44 MRSA strain (**Table 1**).

**Table 1.** MIC determination 21 initially selected 5-sulfonyl-1,3,4-oxadiazole compounds using MRSA.

| ID | R<sub>1</sub> | R<sub>2</sub> | MIC (μg/mL) | ID | R<sub>1</sub> | R<sub>2</sub> | MIC (μg/mL) | ID | R<sub>1</sub> | R<sub>2</sub> | MIC (μg/mL) |
| --- | --- | --- | --- | --- | --- | --- | --- | --- | --- | --- | --- |
| O13 |  | Bn | 23 | O43 |  | *i*-Bu | 42 | O33 |  | Me | 38 |
| O28 |  | Bn | >48 | O48 |  | *i*-Bu | 43 | O44 |  | Me | >44 |
| O30 |  | Bn | 12 | O57 |  | *i*-Bu | 44 | O46 |  | Me | >44 |
| O45 |  | Bn | >46 | O20 |  | *i*-Pr | >46 | O21 |  | *sec*-Bu | 46 |
| O60 |  | Bn | 44 | O59 |  | *i*-Pr | 23 | O32 |  | *sec*-Bu | 44 |
| O36 |  | *i*-Bu | 44 | O5 |  | Me | 40 | O37 |  | *sec*-Bu | >48 |
| O38 |  | *i*-Bu | 42 | O22 |  | Me | 22 | O41 |  | *sec*-Bu | >42 |

From the MIC measurements, several observations were made. First, the lowest MIC was found using compound **O30** (MIC= 25 μM = 12 μg/mL). Second, the R_2_ substituent appears to have a relatively weak correlation with activity as almost every compound containing a benzyl, group at the R_2_ position demonstrated some activity. Third, the addition of electron withdrawing groups at R_1_ had a strong effect on antibacterial activity (**Supplementary Figure S1, Supplementary Table S1**).

Based on the structure of this compound, it was clear that the stereocenter at R_2_/*tert*-butoxycarbonyl (NHBOC) carbon could be an additional variable affecting the activity of these compounds. Therefore, a stereoselective synthesis was developed to produce both the *(R)* and *(S)* forms of compound **O30** (2-methyl-2-propanyl (1-{5-[(3,4-dichlorobenzyl)sulfonyl]-1,3,4-oxadiazol-2-yl}-2-phenylethyl)carbamate; **Scheme 1, Scheme S1, Supplementary Figures S1-10**).

**Scheme 1.**
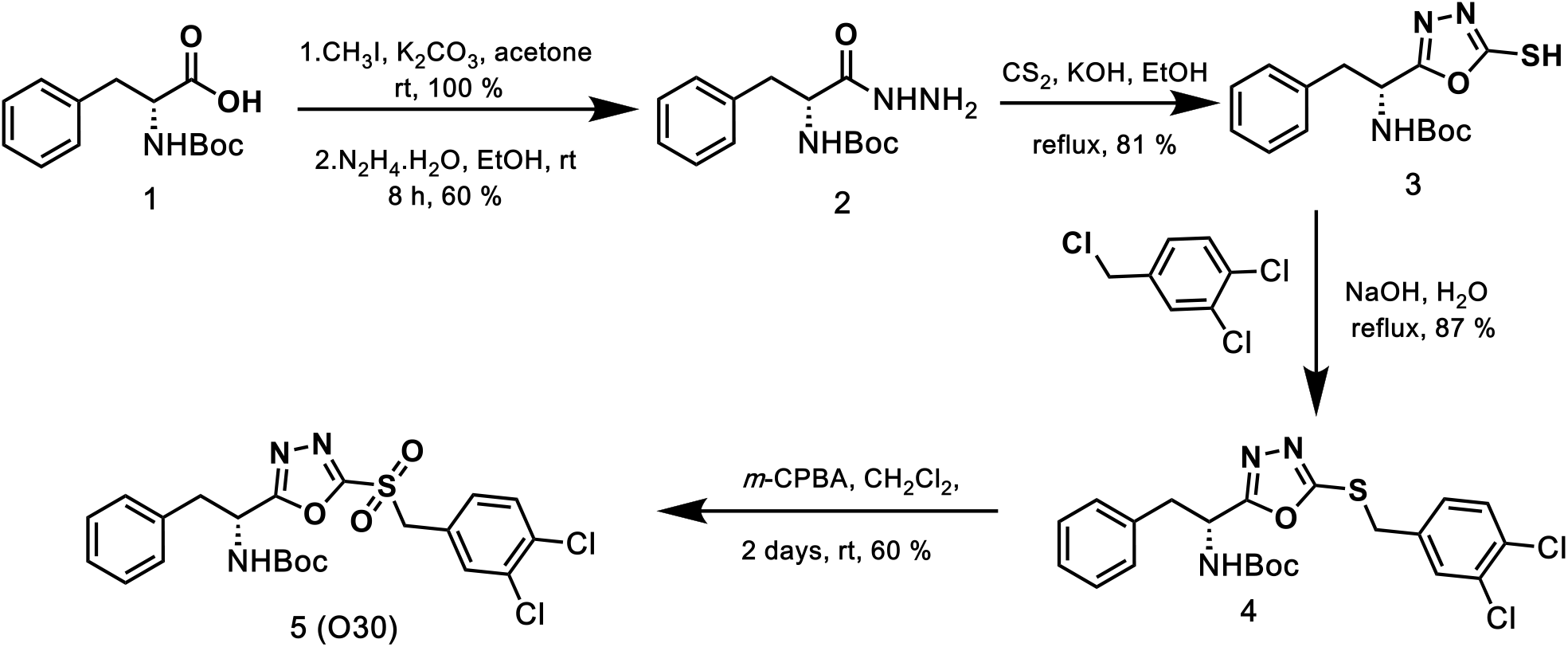
Synthesis of enantiopure *(R)-*O30 (5).

A combination of NMR, mass spectrometry, and Optical rotation was used to confirm the purity of both enantiomers (see Materials and Methods). Subsequently, the overall MIC values for both enantiomer compounds were measured and compared against all in-house laboratory strains of MRSA, including ATCC BAA-44 bioluminescent *S. aureus*, to analyze their inhibition activity (**Table 2a**). This demonstrated that ***(R)*-O30** exhibited 4-8x greater activity compared to its *(S)* enantiomer.

**Table 2.**
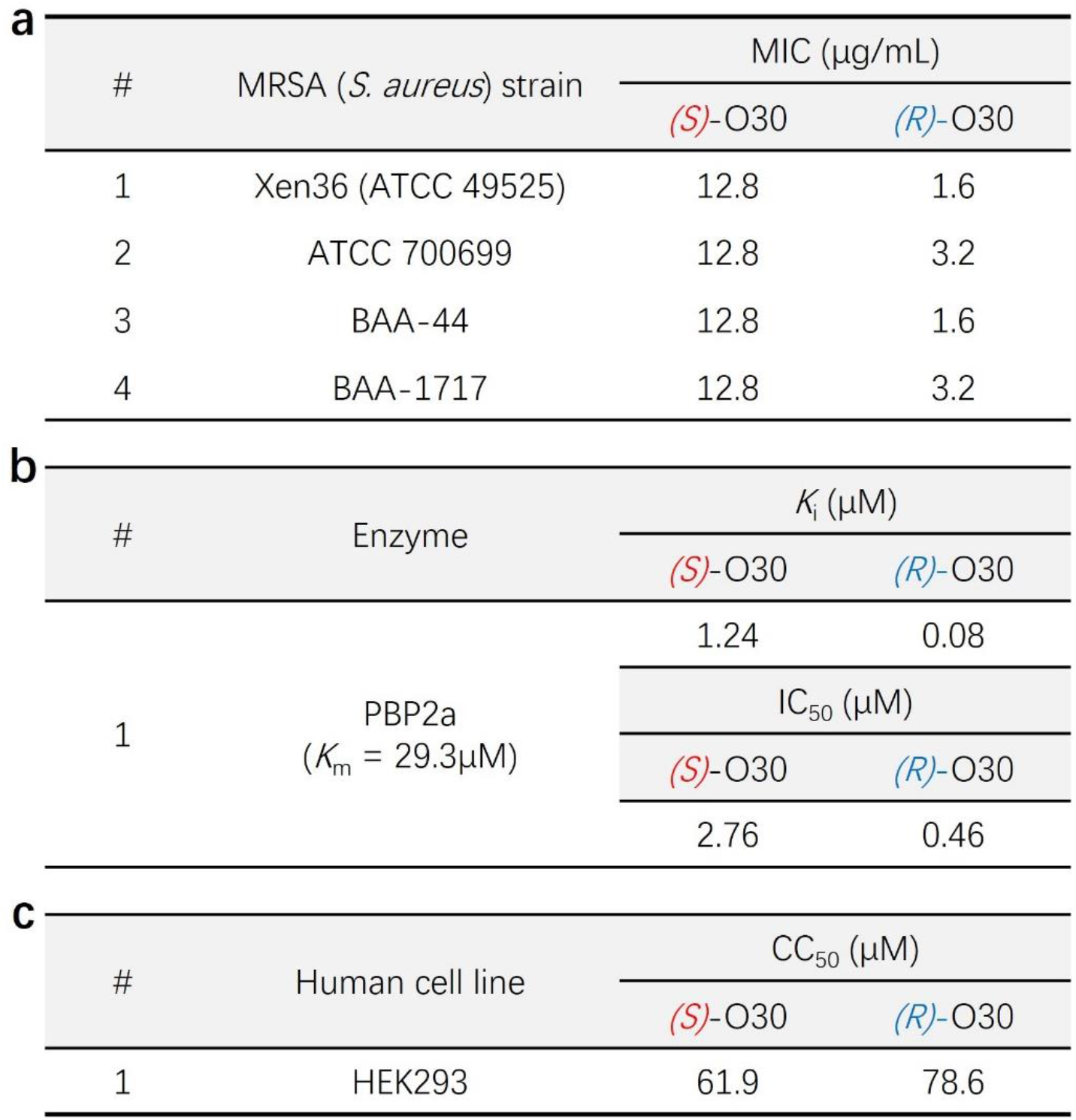
Cell and enzyme level characterization of (*S*)- and (*R*)-O30. **(a)** MIC determination and comparison against various MRSA strains. **(b)** Enzyme inhibition studies against PBP2a using nitrocefin as the substrate. **(c)** Half-maximal cytotoxic concentrations (CC_50_) against human HEK239 cells.

Through repeated experiments, we demonstrated that *(R)*-O30 exhibited significantly increased antibacterial activity compared to the *(S)* enantiomer, representing a ∼4-8x increase in overall cell inhibition activity (**Table 2a**). Given our expectation that PBP2a is the target of these inhibitors, we characterized the enzyme inhibition kinetics of PBP2a with nitrocefin, a chromogenic cephalosporin, as the substrate. The *K*_m_ and *k*_cat_ of PBP2a were determined to be 29.28μM and 259.1 s^-1^, respectively (**Table 2b**). The *K*_i_ and IC_50_ values of both enantiomers were also determined. ()-O30 exhibits a low micromolar inhibitory concentration against PBP2a, whereas *(R)*-O30 is far more potent with a *K*_i_ of 0.08 μM and an IC_50_ of 0.46 μM (**Table 2b**).

Then, to determine whether ***(R)*-O30** could be utilized in further *in vivo* studies, an MTT assay was carried out with human kidney cells (HEK239) to investigate any potential cell toxicity. Both enantiomers exhibited similar half-maximal cytotoxicity concentrations (CC_50_) of 61.9±0.4 μM for ***(S)*-O30** and 73.2±14.8 μM for ***(R)*-O30**, respectively, indicating a cytotoxicity range that is sufficiently acceptable compared to several conventional antibiotics in use. (**Table 2c**).

Based on these promising antibacterial, enzymatic, and cytotoxicity results, further characterization of the *in vitro* activity of ***(R)*-O30** was pursued using 10 clinical MRSA isolates obtained from the group of Dr. Robert Bonomo (VA Northeast Ohio Healthcare System)(**Table 3**). Most (90%) of the clinical isolates exhibit strong resistance to both methicillin and oxacillin. Treatment with ***(R)*-O30** demonstrated highly potent inhibitory activity, with MIC values ranging from 1.8 to 3.6 μg/mL (MIC_50_=3.2 μg/mL, MIC_90_=3.6 μg/mL; **Table 3**) and IC_50_ values in the low micromolar range.

**Table 3.** MIC determination against MRSA Clinical isolate strain using ***(R)*-O30**.

| # | MRSA Strain | Oxacillin resistance | MIC (µg/mL) | IC<sub>50</sub> (µM) |
| --- | --- | --- | --- | --- |
| 1 | SA1622 | R | 1.8 | 1.04 ± 0.95 |
| 2 | SA1623 | S | 3.6 | 1.76 ± 0.53 |
| 3 | SA1624 | R | 1.8 | 1.02 ± 0.88 |
| 4 | SA1626 | R | 3.6 | 1.88 ± 1.00 |
| 5 | SA1627 | R | 1.8 | 1.28 ± 0.84 |
| 6 | SA1628 | R | 3.6 | 2.35 ± 0.92 |
| 7 | SA1629 | R | 3.6 | 1.74 ± 0.91 |
| 8 | SA1630 | R | 3.6 | 1.55 ± 0.92 |
| 9 | SA1633 | R | 1.8 | 2.34 ± 0.87 |
| 10 | SA1637 | R | 3.6 | 1.31 ± 0.89 |

### Application of a PEG-based ointment to a murine skin wound infection model

Based on these promising results, an in vivo study was planned to validate the antibacterial efficacy of this new compound in a more complex biological model. However, this compound is insoluble in water and its current solubility is limited to polar organic solvents such as DMSO. For this reason, two steps were taken. First, a topical PEG-based ointment was prepared with 0.6% ***(R)*-O30** and its in vitro efficacy was validated to ensure there was no measurable reduction in activity **(Figure 1a,b)**.

**Figure 1.**
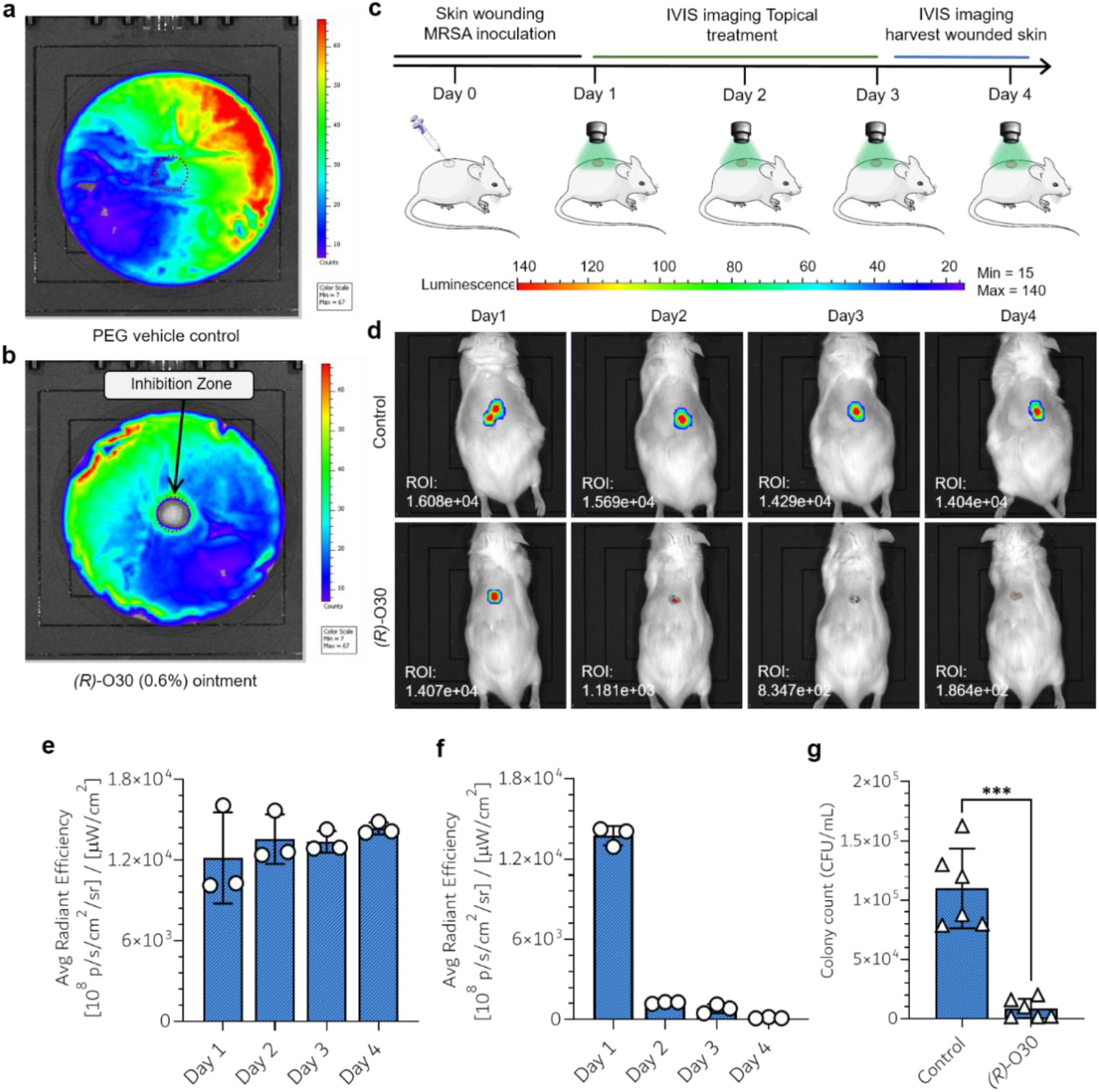
*In vitro* and *in vivo* skin wound infection model tests of PEG-based *(R)*-O30. **(a, b)** Inhibition zone disk diffusion assay comparing PEG vehicle control and 0.6% *(R)*-O30. **c)** Study design; **d)** imaging results; **e)** ROI quantification for mice treated with the vehicle control; **f)** ROI quantification for mice treated with 0.6% *(R)*-O30; **g)** CFU counts from biopsied tissue at the conclusion of the experiment. *** *p < 0*.*001*

By comparing both plates, it could be clearly observed that the PEG-based 0.6% ***(R)*-O30** ointment retained its antibacterial activity, resulting in an obvious 14 mm inhibition zone where a small pellet (50 μL volume pre-liquefied and drop cast-cooled) had been placed (**Fig. 1b)**. In comparison, the vehicle control did not demonstrate any inhibition.

The *in vivo* studied was then carried out using a skin wound infection mouse model. Dorsal biopsies (5 mm) were created on JAX Swiss albino mice and subsequently infected with S. aureus (Xen 36; 40 μL, 5×10^5^ CFU/mL; **Figure 1c**). The extent of infections (and treatment) was monitored using the IVIS Lumina system in the bioluminescence imaging mode. Treatments were applied once daily for three days and images were taken prior to each treatment. From these images and from further region of interest (ROI) quantification, it could be clearly observed, in real time, that the ***(R)*-O30** topical ointment quickly and significantly decreased bacterial burden **(Figure 1d-f)**. After one treatment, the image (day 2) demonstrates a more than 10x decrease in bacterial bioluminescence as compared to the control. However, to quantify the decrease in bacterial concentration, the infected tissue with an additional 1 mm margin (6 mm biopsy) was biopsied, homogenized, and plated to count the number of colonies forming units (CFU). After 18 hours, a clear and significant (p<0.001) difference could be observed between the ***(R)*-O30** and vehicle control treated mice (**Figure 1g**). These promising results suggest that formulations containing ***(R)*-O30** can effectively combat full-depth excisional wounds, which act as a model for severe skin and soft tissue infections (SSTI).

### Observing allosteric inhibition of PBP2a via covalent binding

Given the observed efficacy of *(R)*-O30 against various MRSA strains and the known structural active site similarities between PBPs and β-lactamases, we hypothesized that PBP2a was a likely bacterial target. Sequential LC-MS/MS studies and computer modelling were therefore used to investigate the potential binding mechanism between *(R)*-O30 and the PBP2a protein (one of the main drivers of drug resistance in MRSA; **Figure 2,3**).

**Figure 2.**
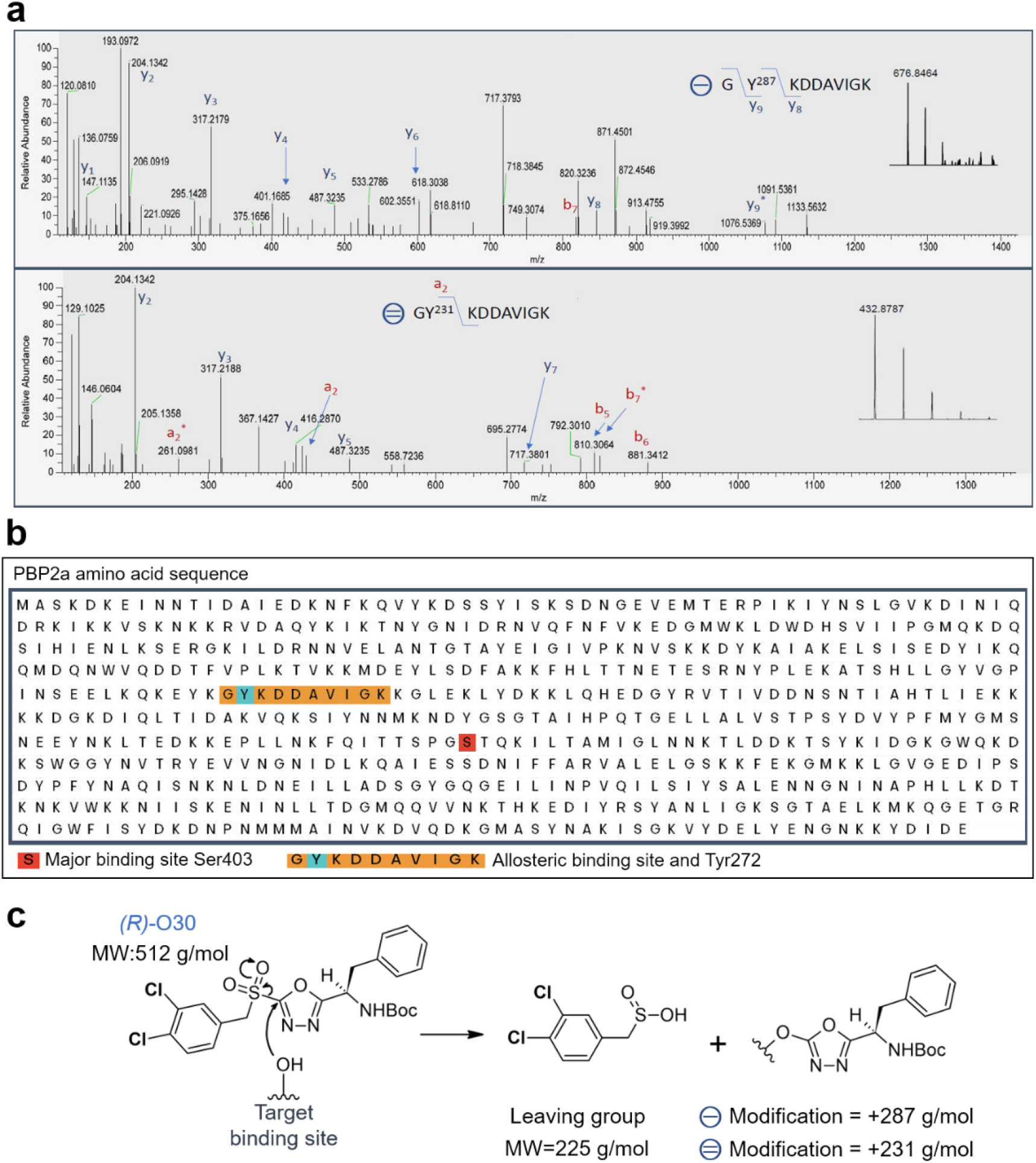
Combined mass spectrometry and sequence analysis confirm allosteric binding via oxadiazole-ring substitution. **a)** LC-MS/MS analyses indicating 287 Da and 231 Da tyrosine residue modification. **b)** PBP2a amino acid sequence with major binding site (red) and allosteric site (orange) with modified residue (cyan). **c)** Proposed nucleophilic aromatic substitution (S_N_Ar) reaction mechanism based on the observed peptide modifications.

Our LC-MS/MS analysis, conducted after treating PBP2a with *(R)*-O30, revealed the formation of a covalent bond at tyrosine (Y) 223/272 located at an allosteric site of PBP2a. This increases the mass of this tyrosine residue by either 231 g/mol or 287 g/mol, depending on molecular fragmentation (**Figure 2a,b)**. This finding supports a nucleophilic aromatic substitution (S_N_Ar) reaction as the mechanism for the covalent binding at the allosteric site (**Fig. 2c**). In this mechanism, the sulfonyl group leaves and this may in part explain our previous observation that electron withdrawing groups, such as chlorines, increased the apparent antimicrobial efficacy of these compounds (**Table 1**).

To further understand which molecular interactions may assist the association and docking of (*R*)-O30 into the allosteric site, a predictive model for the covalent binding between *(R)*-O30 and tyrosine 223/272 was constructed based on the analyzed LC-MS/MS data (**Figure 3a,b**). From this model, several important interactions are observed that may be crucial for enabling the S_N_Ar reaction (**Figure 3c)**. These include hydrogen bonding with K273 and H293 and a likely cation-π interaction with between the aromatic benzyl group and K319. These results also help to explain our previous MIC and single-dose assay observations, wherein compounds that had 3,4-dichlorobenzyl groups as R_1_ demonstrated the following trend in activity according to their R_2_ groups: benzyl > isopropyl > methyl >> *sec*-butyl (**Table 1, Supplementary Table S1)**. We speculate that the smaller isopropyl and methyl substituents neither enhance nor hinder activity, which still relies on docking via H-bond interactions and covalent inhibition via S_N_Ar reaction. Further work is needed with careful design of alternative R_2_ groups to precisely identify a best fitting compound.

**Figure 3.**
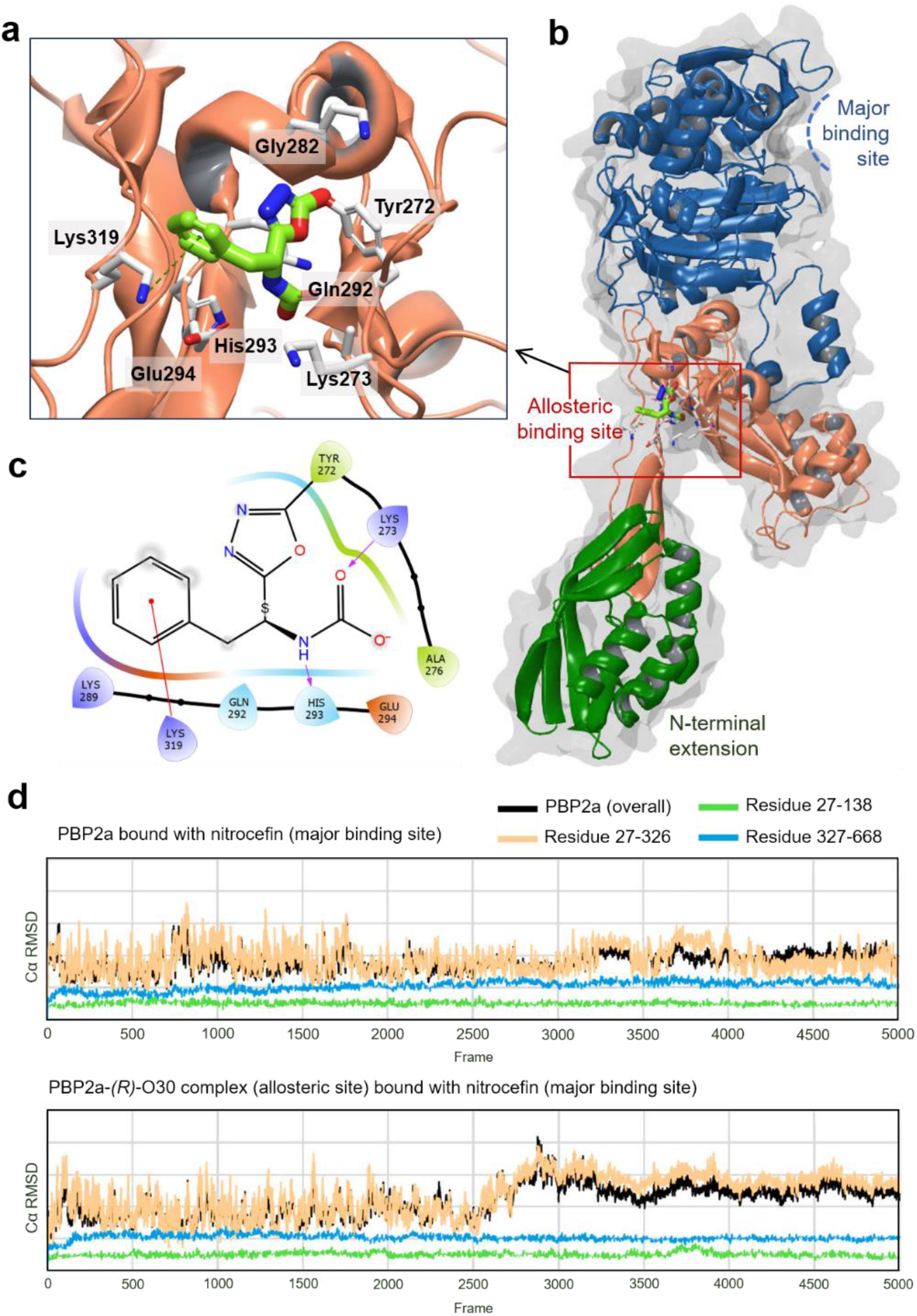
Covalent docking and dynamic computational modeling. **(a)** Covalent inhibition model of *(R)-*O30 at the highlighted allosteric sight. (b) Crystal structure of PBP2a (PDB: 1VQQ) indicating the active site (blue), the allosteric site (orange), and the N-terminal extension (green). (c) Ligand interaction diagram for *(R)*-O30 at the PBP2a allosteric site. (d) Long-term molecular dynamic simulations of PBP2a: nitrocefin without (*R*)-O30 (bottom) and with *(R)*-O30 at the allosteric site (top).

According to several research reports, the activity of the PBP2a protein is known to be regulated by allosteric sites distinct from the active site where cell wall cross-linking occurs^28,29^. These initial reports detail the importance of peptidoglycan activators binding at this key allosteric site located in Lobe 2^30^. This binding results in a cascade of salt bridge modifications that ultimately open the otherwise closed active site, which is located more than 60 Å away^30^. This same mechanism has observed with MRSA-active cephalosporins such as ceftaroline or ceftobiprole; both have been observed to dock at the Lobe 2 allosteric position as well as the active site^28,29^.

To better understand this process with our new inhibitor, protein models of PBP2a with and without *(R)*-O30 were subjected to long-term molecular dynamic simulations using the Schrodinger package, Desmond. Doing so, the spatiotemporal and dynamic structural changes of the major active site, the allosteric site, and the N-terminal extension were examined (**Figure 3d**). These experiments were conducted with nitrocefin in the active site and the *(R)*-O30-free and *(R)*-O30-bound models were compared. Our results agree very well with previous studies on MRSA-active cephalosporins, demonstrating that the active site of the *(R)*-O30-bound PBP2a dynamically changes conformation (**Figure 3d**). The active site of the *(R)*-O30-free PBP2a, on the other hand, remains unchanged over the course of the simulation.

Upon closer inspection of the allosteric site and which residues are involved, we observe several common residues involved in the docking of *(R)*-O30 and other ligands in Lobe 2: K319, K273, and H293^29^. These all confirm that *(R)*-O30 docks are in the same key, allosteric site. However, unlike other ligands, LC-MS/MS data confirms that *(R)*-O30 binds covalently and therefore irreversibly locks PBP2a into its new confirmation (**Figure 2**).

However, given that *(R)*-O30 was not found to be bound to the active site of PBP2a, we speculate that the induced PBP2a conformational changes result in either a closed or an *inactive* semi-open state where peptidoglycan fragments are either still restricted from entering the active site or are able to enter but unable to adopt the correct configuration for an efficient linking reaction. Given the fact no *(R)*-O30 was found in the active site, we tend towards the first option in which the salt-bridge cascade has been disrupted and the active site either remains closed or is not sufficiently open to permit entry to small molecules.

We therefore believe that our *(R)*-O30 may be a template for superior PBP2a effectors/inhibitors or combination therapy drugs given that binds irreversibly and it does not mimic a β-lactam or a nascent peptidoglycan^28,31^.

## Conclusion

Through combined *in silico* modeling, in vitro, and in vivo approaches, we have demonstrated that 5-dichlorobenzylsulfonyl-1,3,4-oxadiazoles are a promising class of compounds for the treatment of MRSA infections. They exhibit low toxicity, high activity, and a new mechanism of action based on allosteric modulation of PBP2a—a key target for MRSA active agents. Our future studies will investigate the potential for synergism between these agents and other MRSA active agents as well as search for derivatives with increased activity. Generally, we believe that by targeting allosteric sties of drug resistant enzymes will offer unique opportunities for novel drug discovery with increased efficacy and potentially decreased side effects.

## Materials and Methods

### Chemical synthesis

All reactions were carried out in oven-dried glassware (unless water was present in the reaction mixture) with magnetic stirring under a positive pressure of argon unless otherwise indicated. ACS reagent grade solvents were used. Reactions were monitored by thin layer chromatography (TLC) carried out on 250 μm Merck silica gel plates (60 F254) containing a fluorescent indicator (254 nm). TLC plates were visualized under an iodine chamber or UV lamp before treatment with the cerium ammonium molybdate stain and development with heat. Flash column chromatography was performed using Silicycle SiliaFlash P60 silica gel (60 Å pore size, 40 – 63 μm particle size, 230 – 400 mesh) and ACS reagent grade solvents. Melting points were determined with a Mel-Temp Digital Melting Point Apparatus. Optical rotation was measured using LAXCO POL-200 Series Automatic Polarimeter. Infrared spectra were recorded on a Bruker Tensor 27 FT-IR spectrometer and reported as wavenumber (cm^−1^). High-resolution mass spectra (HRMS) were recorded using Thermo Scientific Exactive Plus Orbitrap Mass Spectrometer. ^1^H NMR and ^13^C NMR spectra were recorded in (CD_3_)_2_SO or CDCl_3_ on a Bruker Av 400 MHz NMR-spectrometer operating at 400 MHz (^1^H), 101 MHz(^13^C) and Agilent 500 MHz NMR-spectrometer operating at 500 MHz (^1^H) and 125 MHz (^13^C). Chemical shifts (δ) are reported in ppm and are referenced to the chemical shift of the residual solvent proton(s) present in dimethyl sulfoxide δ[ppm] = ((CH_3_)_2_SO) = 2.50 ppm for the ^1^H NMR spectra and δ[ppm] = ((CD_3_)_2_SO) = 39.52 ppm for the ^13^C NMR spectra, or chloroform δ[ppm] = (CDCl_3_) = 7.26 ppm for the ^1^H NMR spectra and δ[ppm] = (CDCl_3_) = 77.16 ppm for the ^13^C NMR spectra. The corresponding peak multiplicities are abbreviated as follows: s = singlet, d = doublet, t = triplet, q = quartet, and m = multiplet. Coupling constant values were extracted assuming first-order coupling and shown in hertz (Hz).

### D-Boc-phenylalanine methyl ester (1a)

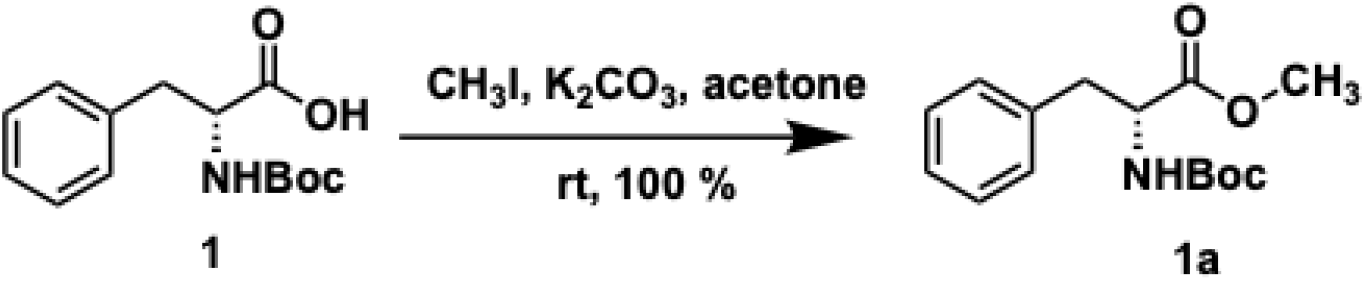

To the solution of carboxylic acid **1** (18.85 mmol, 5.00 g) in acetone (0.60 M, 30 mL) in a 100 mL RB flask, K_2_CO_3_ (2.00 equiv., 37.70 mmol, 5.21 g) and CH_3_I (3.00 equiv., 56.54 mmol, 3.50 mL) were added, and the mixture was stirred overnight. The solvent was removed under reduced pressure, diluted with NH_4_Cl (15 mL), extracted with EtOAc (2 X 20 mL), washed with brine (15 mL), dried over anhydrous Na_2_SO_4,_ and concentrated under reduced pressure to give a pure colorless oil, methyl ester **1a** (5.37 g,19.22 mmol, 100 %).

R_f_: 0.62 (40 % EtOAc in hexanes)

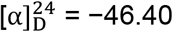 (c = 1.0 in CHCl_3_)

IR (neat): v_max =_ 3438, 3366, 3029, 2978, 2932, 1744, 1712, 1496, 1365, 1162 cm^-1^

^1^H NMR (500 MHz, CDCl_3_) δ 7.51 – 7.20 (m, 5H), 5.18 (d, J = 8.6 Hz, 1H), 4.80 – 4.65 (m, 1H), 3.82 (s, 3H), 3.29 – 3.08 (m, 2H), 1.54 (s, 10H)

^13^C NMR (101 MHz, CDCl_3_) δ 172.32, 155.07, 136.05, 129.24, 128.47, 126.94, 79.76, 54.42, 52.11, 38.24

HRMS-ESI (m/z): calcd. for C_15_H_21_NNaO_4_^+^ ([M + Na] ^+^): 302.1363, found: 302.1363.

### N-Boc-D- phenylalanine hydrazide (2)^33^

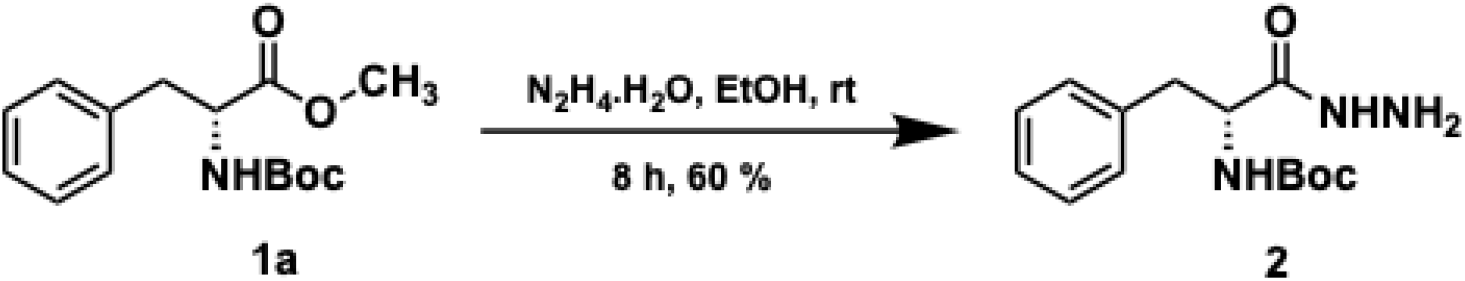

In a 100 mL RB flask, the solution of methyl ester **1a** (15.60 mmol, 4.37 g), ethanol (0.60 M, 26 mL), and N_2_H_4_.H_2_O (3.00 equiv., 47.00 mmol, 2.40 mL) was prepared and was stirred for 8 hours until TLC analysis indicated complete consumption of starting material. The solvent was removed under reduced pressure to give the white residue which on recrystallization in EtOAc/hexanes afforded white powder hydrazide **2** (2.63 g,9.42 mmol, 60%).

MP: 126 –127 ^o^C

R_f_: 0.05 -0.15 (40 % EtOAc in hexanes)

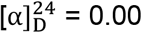 (c = 1.0 in CHCl_3_)

IR (neat): v_max_ = 3324, 2988, 1690, 1658, 1520, 1444, 1242, 1164 cm^-1^

^1^H NMR (500 MHz, CDCl_3_) δ 7.65 (s, 1H), 7.33 – 7.13 (m, 5H), 5.27 (d, J = 8.8 Hz, 1H), 4.36 (d, J = 7.8 Hz, 1H), 3.84 (s, 2H), 3.19 – 2.88 (m, 2H), 1.37 (s, 9H)

^13^C NMR (101 MHz, CDCl_3_) δ 172.06, 136.46, 129.41, 128.92, 127.27, 80.63, 54.83, 38.54, 28.39

HRMS-ESI (m/z): calcd. for C_14_H_21_N_3_NaO_3_^+^ ([M + Na] ^+^): 302.1475, found: 302.1475.

### tert-butyl N-[(1R)-1-(4,5-dihydro-5-thioxo-1,3,4-oxadiazol-2-yl)-2-phenylethyl] carbamate (3)

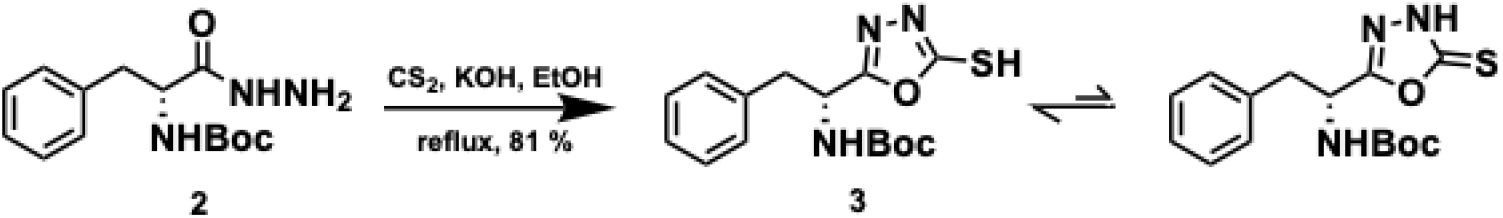

To the solution of hydrazide **2** (13.96 mmol, 3.90 g) in dry ethanol (0.18 M, 75 mL) in a 250 mL RB flask, KOH (1.20 equiv., 16.76 mmol, 940 mg) was added under argon at room temperature and was stirred for an hour. A solution of CS_2_ (1.20 equiv., 16.76 mmol, 1.01 mL) was prepared in dry ethanol (5 mL) and slowly added to the above solution and heated to reflux overnight. The solvent was evaporated and CH_3_COOH (2 mL) was added to get the white precipitate which was recrystallized in EtOAc/hexanes to give **3** (2.47 g, 7.69 mmol, 55 %). However, on flash chromatography at 20 % EtOAc in hexanes, **3** can be obtained up to 81 %. Note: Anhydrous ethanol was prepared in oven-dried RB flask with 4 Å molecular sieves under argon.

MP: 145 –147 ^o^C

R_f_: 0.35 (40 % EtOAc in hexanes)

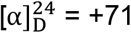 (c = 1.0 in CHCl_3_)

IR (neat): v_max_ =3339, 3172, 2987,2965, 2934, 1689, 1616, 1523, 1470, 1165 cm^-1^

^1^H NMR (400 MHz, CDCl_3_) δ 7.38 – 7.04 (m, 5H), 5.30 – 4.74 (m, 2H), 3.18 (qd, J = 14.1, 6.6 Hz, 2H), 1.36 (d, J = 25.6 Hz, 9H)

^13^C NMR (101 MHz, CDCl_3_) δ 178.66, 162.89, 155.06, 134.63, 129.28, 128.91, 127.52, 81.39, 48.48, 38.49, 28.27

HRMS-ESI (m/z): calcd. for C_15_H_19_N_3_NaO_3_S^+^ ([M + Na] ^+^): 344.1038, found: 344.1039.

### tert-butyl N-[(1R)-1-[5-[[(3,4-dichlorophenyl) methyl] thio]-1,3,4-oxadiazol-2-yl]-2-phenylethyl] carbamate (4)

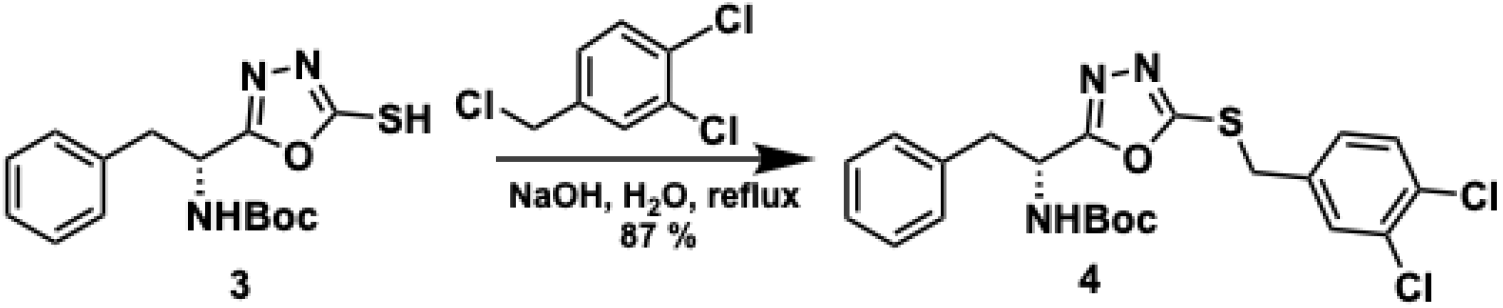

The mixture of thio-oxadiazole **3** (10.39 mmol, 2.30 g) and deionized water (0.92 M, 20 mL) was prepared in a 100 mL RB flask. Then, NaOH (1.20 equiv., 12.47 mmol, 500 mg) and 3,4-dichlorobenzyl chloride (1.50 equiv., 15.59 mmol, 2.16 mL) were added and heated to reflux for 36 hours. The reaction mixture was cooled and then diluted with NH_4_Cl (10 mL) and 20 mL EtOAc. The layers were separated, and the aqueous layer was extracted with two additional portions of EtOAc (20 mL each). The combined organics were washed with brine (10 mL) before drying over anhydrous Na_2_SO_4_. The solvent was removed under reduced pressure to afford benzylated product **4** (2.98 g, 6.20 mmol, 87 %).

MP: 112 –115 ^o^C

R_f_: 0.67 (40 % EtOAc in hexanes)

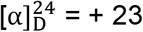 (c = 0.75 in CHCl_3_)

IR (neat): v_max_ =3365, 1691, 1519, 1478, 1247, 1156, 876, 725 cm^-1^

^1^H NMR (500 MHz, CDCl_3_) δ 7.60 – 7.02 (m, 9H), 5.26 (d, J = 6.7 Hz, 1H), 5.05 (d, J = 9.8 Hz, 1H), 4.35 (s, 2H), 3.35 – 3.04 (m, 2H), 1.41 (s, 9H)

^13^C NMR (101 MHz, CDCl_3_) δ 167.49, 163.66, 154.77, 136.00, 135.23, 132.76, 132.30, 130.99, 130.68, 129.31, 128.71, 128.53, 127.28, 80.58, 48.33, 39.64, 35.35, 28.24

HRMS-ESI (m/z): calcd. for C_22_H_23_Cl_2_N_3_NaO_3_S^+^ ([M + Na] ^+^): 502.0732, found: 502.0729.

### tert-butyl N-[(1R)-1-[5-[[(3,4-dichlorophenyl) methyl] sulfonyl]-1,3,4-oxadiazol-2-yl]-2-phenylethyl] carbamate (5)^34^

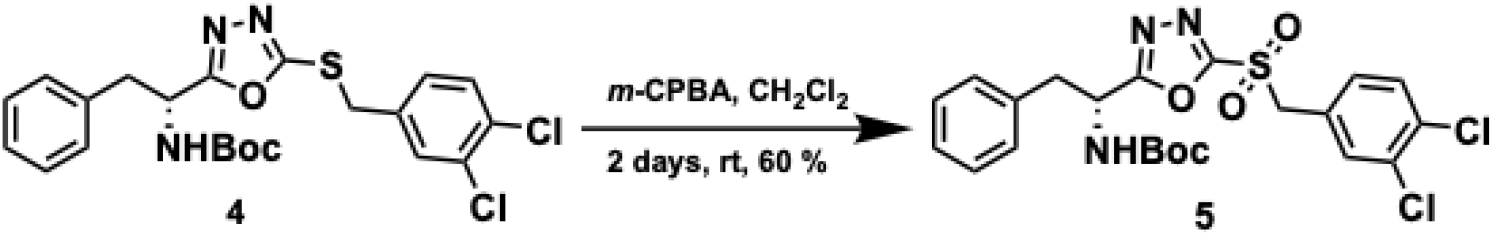

To the solution of sulfide **5** (0.78 mmol, 377 mg) in CH_2_Cl_2_ (0.05 M, 15 mL), 70 % *m*-CPBA (4.00 equiv., 3.12 mmol, 700 mg) was added at 0 ^o^C and stirred for 2 days at room temperature. The reaction mixture was diluted with 2 M NaOH (10 mL), extracted with CH_2_Cl_2_ (2 X 20 mL), washed with brine (10 mL), dried over anhydrous Na_2_SO_4,_ and concentrated under reduced pressure to get crude (411 mg) which on recrystallization in methanol gave pure sulfone **5** (213 mg, 0.42 mmol, 60 %).

MP: 132 –134 ^o^C

R_f_: 0.70 (40 % EtOAc in hexanes)

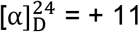 (c = 0.50 in CHCl_3_)

IR (neat): v_max_ = 3336, 1687, 1521, 1468, 1350, 1315, 1253, 1167, 1034 cm^-1^

^1^H NMR (400 MHz, CDCl_3_) δ 7.49 (d, J = 2.2 Hz, 1H), 7.45 (d, J = 8.2 Hz, 1H), 7.32 – 7.26 (m, 3H), 7.20 – 7.05 (m, 3H), 5.38 – 5.25 (m, 1H), 5.03 (d, J = 9.3 Hz, 1H), 4.67 (s, 2H), 3.31 – 3.13 (m, 2H), 1.40 (s, 9H)

^13^C NMR (101 MHz, CDCl_3_) δ 169.22, 161.52, 154.77, 134.86, 134.56, 133.68, 133.21, 131.31, 130.59, 129.27, 129.12, 127.81, 124.81, 81.22, 60.52, 48.83, 39.56, 28.30

HRMS-ESI (m/z): calcd. for C_22_H_23_Cl_2_N_3_NaO_5_S^+^ ([M + Na] ^+^): 534.0631, found: 534.0628.

### Expression and Purification of Staphylococcus aureus PBP2a

A gene fragment containing PBP2a (residues 23–668) from the Staphylococcus aureus strain was amplified using PCR and subsequently cloned into a PET28a plasmid via NdeI/XhoI restriction digestion, resulting in a PET28a-PBP2a construct. This construct features an octa-histidine (6x His) tag at the C-terminus of PBP2a. The resulting plasmid was then transformed into E. coli BL21(DE3) cells (Novagen) for protein expression. The bacterial cells were cultured in Luria broth (LB), supplemented with an antibiotic (100 μg/ml ampicillin), and incubated at 37 °C until the optical density of the culture reached 0.6 at a wavelength of 600 nm (OD600). The expression of PBP2a was induced with 0.1 mM isopropyl-β-D-thiogalactopyranoside (IPTG) and incubated at 25 °C for 16 hours. Native SauPBP2a was expressed as a soluble protein and purified as described with some modifications^[1]^ .

The cultures were harvested by centrifugation, and the cell pellets were resuspended in a purification buffer (20 mM Tris-HCl pH 7.5 and 150 mM NaCl), supplemented with 0.1 mM phenylmethylsulphonyl fluoride (PMSF, Sigma-Aldrich). The cells were lysed by passing them three times through a cell disrupter (ATS Engineering Ltd), and cell debris was removed by centrifugation at 18,000 x g for 45 minutes at 4 °C. The solubilized protein suspension was ultracentrifuged at 100,000 x g for 1 hour before being loaded onto a 5 ml HisTrap HP column (GE HealthCare). The column was washed with a purification buffer supplemented with 20 mM imidazole, and the bound protein was eluted with a purification buffer supplemented with 250 mM imidazole. The protein eluted from the HisTrap HP column was further purified by size-exclusion chromatography using a Superdex 200 Increase 10/300 column (GE Healthcare) equilibrated in 20 mM Tris-HCl pH 7.5, 150 mM NaCl. The purity of the protein fractions was analyzed by SDS-PAGE. Fractions with the highest purity were collected and concentrated to 5 mg/ml for further study.

### Biochemical assays

Nitrocefin was used as the substrate as this is a chromogenic cephalosporin. The acrtivity of purified PBP2a was measured spectrophotometrically (Cytation 3, BioTek) in potassium phosphate buffer (PBS, pH = 7.4, prepared according to published recipe). Nitrocefin concentrations ranging from 50 μM to 200 μM were utilized. The formation of hydrolyzed product was measured at 486 nm at intervals of 10 s for at least 30 min until a plateau in product formation was observed. The *K*_*m*_ and *k*_*cat*_ values were determined by plotting initial velocity measurements, followed by fitting with a non-linear regression Michaelis-Menten kinetic model in GraphPad Prism 10.

The inhibitor constants (*k*_*i*_) were determined following a similar approach, where the inhibitor concentrations ranged from 8-128 μM and nitrocefin ranged from 50-200 M. Each inhibitor was pre-incubated with the PBP2a enzyme for 15 min at room temperature, and then the absorbance was measured at 486 nm. The results were evaluated using GraphPad Prism 10 andf the previously determined k_m_ value.

### Disc diffusion assay

Overnight cultured bacteria were diluted to 0.5 McFarland Standard. Kirby-Bauer discs containing either 0.6 % (*R*)-O30 or delivery vehicle were added to the inoculated Müller-Hinton Agar (MHA) plates and incubated overnight at 37 °C.^43-45^The following day photos and luminescence images were recorded to determine the inhibition zones around each disc.

### Single Dose inhibition assay

Bacteria were diluted to 0.5 McFarland Standard in MHB2. A growth control containing only bacteria and a negative control containing only MHB were added. Test wells were treated with either 50 μM meropenem or amoxicillin. Each set was repeated in triplicate and the final viability was determined in reference to the growth control.

### Luminescence Imaging

Luminescent images were obtained using the IVIS^®^ Lumina XRMS Series III (PerkinElmer) in bioluminescence imaging mode. Exposure times were automatically determined by the LivingImage Software with medium binning. Mice were kept under anesthesia using the attached XGI-8 Gas Anesthesia System from Caliper LifeSciences.

### Minimum inhibitory concentration (MIC) assay

MIC assays were carried out following established CLSI guidelines using MHB-2 and bacterial concentrations equivalent to 0.5 McFarland Standard.^46,47^ The MIC was reported as the concentration at which no growth was observed.

### In vivo treatment of murine skin wound model

Xen 36 *S. aureus* was grown overnight in MHB-2 broth. The overnight culture was then centrifuged, the media removed, and the pellet was resuspended in PBS. This suspension was later used to inoculate the mice. Anesthesia was induced with 3-5% isoflurane and maintained with 2-3% isoflurane. Mice were then shaved, and the dorsal skin was scrubbed with povidone and washed away with 70% ethanol. A clear plastic template with a square 1 cm^2^ cutout was used to demarcate the corners of the planned needle-scratch grid. A 25G, 1½ needle was then used to carefully create the abrasive wound, with caution being paid not to deeply lacerate (See supplementary note S1 for additional details). Having created the two sets of wounds, Xen 36 (40 μl, 5×10^5^ CFU/mL) was added via pipette to each wound. IVIS images were taken after 24 hours, and this was determined to be the starting point of the experiment (t=0). Treatments were for 3 days, once per day, given while animals were anesthetized by adding either 0.6% DO30 or delivery vehicle only. IVIS images were taken once per day prior to any planned treatment. Following the final treatment on day 3, IVIS images were taken for an additional day.

### Docking and Molecular Dynamics studies

Modeling and docking studies were carried out with the Maestro Schrodinger 2022-2 software package.^35^ Computational studies were based on NMR and X-ray structures (PDB: 1VQQ). Proteins were prepared through the standard workflow, which includes adding missing sidechains and hydrogens as well as energy minimization using the OPLS-2005 force field. Glide docking was utilized to generate docking scores and compare models.^36,37^

MD simulations were performed for PBP2a in complex with the structurally solved inhibitors. Each system was solvated in a cubic box with explicit TIP3P water and nutrient ions consisting a solvent buffer region from the 10 Å edge of the complex. A 100 ns simulation was carried out for the docked model using DESMOND (Research DES. Desmond Molecular Dynamics System. NY, USA 2008) with OPLS-AA 2005 force field under the isobaric-isothermal (NPT) condition at 300 K. The stability of the simulation was assessed by monitoring the CαRMSD (Root-mean-square deviation of α–carbon) with respect to the minimized starting structure.

## Supporting information

R-O30-MRSA_BioRxiv_Shin_SI

## Acknowledgments

Research support for Woo Shik Shin was provided by the Northeast Ohio Medical University, College of Pharmacy. The Northeast Ohio Medical University Department of Pharmaceutical Sciences provided all the necessary research resources.

## Author Contributions

B.M.B. and R.K. provided the overall concept for the compound modification and were responsible for its synthesis. B.M.B., T.H., and A.C. conducted the *in-vitro* and *in-vivo* experiments. C.S. performed the synthesis of the protein enzyme. T.K. and Y.Y.S. conducted and analyzed the *in-silico* MD simulations. R.B. analyzed the results related to the clinical isolates. J.D.M. managed and supervised the compound modification and synthesis, as well as the data analysis. B.M.B., J.D.M., and W.S.S. wrote the manuscript. W.S.S. supervised the work. All authors discussed the results and commented on the manuscript.

## Data Availability Statement

The datasets generated during and/or analyzed during the current study are available from the corresponding author upon reasonable request.

