## Supplementary material for "Development of non-β-Lactam covalent allosteric inhibitors targeting PBP2a in Methicillin-Resistant *Staphylococcus aureus*": R-O30-MRSA_BioRxiv_Shin_SI: R-O30-MRSA_BioRxiv_Shin_SI_final.docx

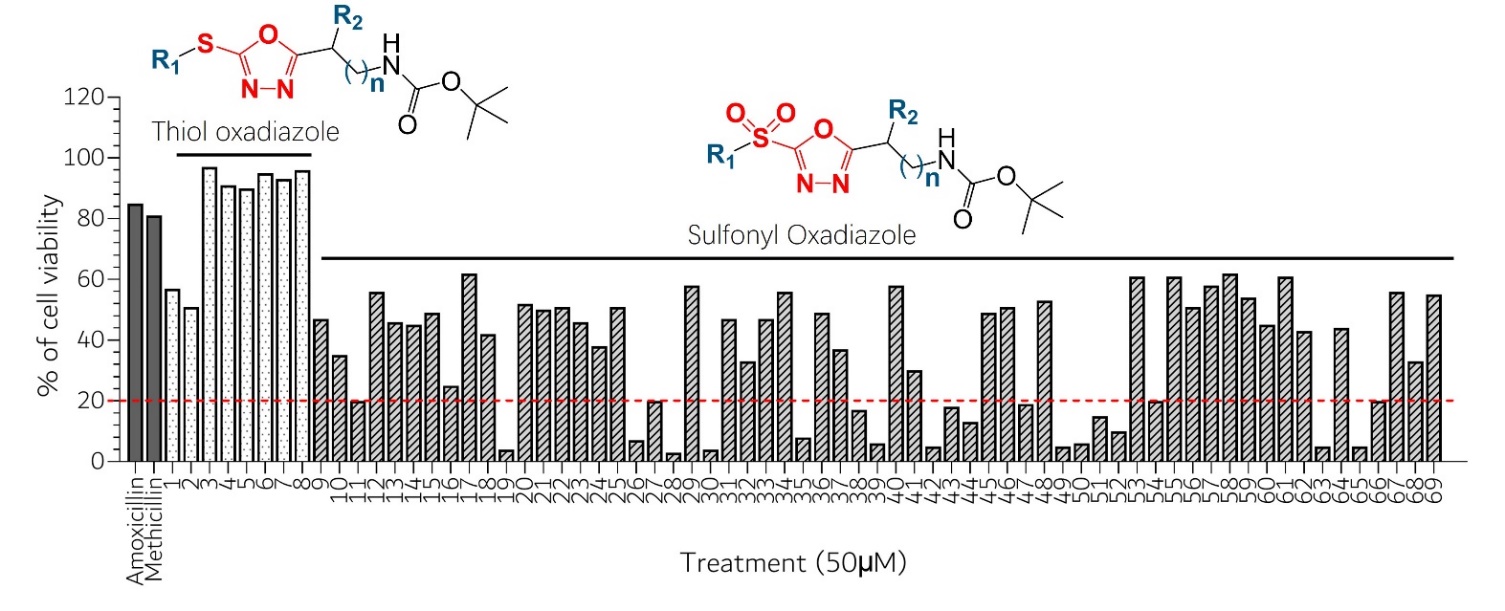

### Figure S1. Initial 77 compound single-dose activity screening against MRSA. Structure-Activity Relationship among the 1,3,4-Oxadiazoles series with single-dose cell-based assay against MRSA pathogen General Chemical Scaffold and single-dose activity screening against MRSA. Structure-Activity Relationship among the 1,3,4-Oxadiazoles series with single-dose cell-based assay against MRSA pathogen c) MIC determination for the MRSA strain of the 21 initially selected 2-sulfonyl-1,3,4-oxadiazole compounds.

**Table S1.** Structure-Activity Relationship among the 1,2,4-Oxadiazoles series with single-dose cell based assay against MRSA pathogen**.**

| Compound | 2' | R_1_ | R_2_ | n | %viability |
| --- | --- | --- | --- | --- | --- |
| Amoxicillin | **-** | - | - | - | 85 |
| Oxacillin | **-** | - | - | - | 81 |
| 1 | **–S–** | 3,4-dichlorobenzyl | - | 1 | 57 |
| 2 | **–S–** | 2,4-dichlorobenzyl | - | 1 | 51 |
| 3 | **–S–** | 2,6-dichlorobenzyl | - | 1 | 97 |
| 4 | **–S–** | 2-chlorobenzyl | - | 1 | 91 |
| 5 | **–S–** | 2-fluoro-6-chlorobenzyl | - | 1 | 90 |
| 6 | **–S–** | 3-chlorobenzyl | - | 1 | 95 |
| 7 | **–S–** | 4-chlorobenzyl | - | 1 | 93 |
| 8 | **–S–** | Benzyl | - | 1 | 96 |
| 9 | **–SO_2_–** | 2-methylbenzyl | methyl | 0 | 47 |
| 10 | **–SO_2_–** | 4-fluorobenzyl | methyl | 0 | 35 |
| 11 | **–SO_2_–** | 4-chlorobenzyl | methyl | 0 | 20 |
| 12 | **–SO_2_–** | 2-chlorobenzyl | methyl | 0 | 56 |
| 13 | **–SO_2_–** | 2-fluoro-6-chlorobenzyl | methyl | 0 | 46 |
| 14 | **–SO_2_–** | 2-fluorobenzyl | *sec*-butyl | 0 | 45 |
| 15 | **–SO_2_–** | 4-fluorobenzyl | - | 1 | 49 |
| 16 | **–SO_2_–** | 4-chlorobenzyl | - | 1 | 25 |
| 17 | **–SO_2_–** | 2-chlorobenzyl | - | 1 | 62 |
| 18 | **–SO_2_–** | 2,5-dimethylbenzyl | - | 1 | 42 |
| 19 | **–SO_2_–** | 2-methylbenzyl | benzyl | 0 | 4 |
| 20 | **–SO_2_–** | 3-methylbenzyl | benzyl | 0 | 52 |
| 21 | **–SO_2_–** | 2-fluoro-6-chlorobenzyl | isobutyl | 0 | 50 |
| 22 | **–SO_2_–** | 2,6-dichlorobenzyl | isobutyl | 0 | 51 |
| 23 | **–SO_2_–** | 2-fluorobenzyl | isopropyl | 0 | 46 |
| 24 | **–SO_2_–** | 4-chlorobenzyl | isopropyl | 0 | 38 |
| 25 | **–SO_2_–** | 2-chlorobenzyl | isopropyl | 0 | 51 |
| 26 | **–SO_2_–** | 2,6-dichlorobenzyl | isopropyl | 0 | 7 |
| 27 | **–SO_2_–** | 2-fluoro-6-chlorobenzyl | *sec-*butyl | 0 | 20 |
| 28 | **–SO_2_–** | 2,6-dichlorobenzyl | methyl | 0 | 3 |
| 29 | **–SO_2_–** | 2,4-dichlorobenzyl | isopropyl | 0 | 58 |
| 30 | **–SO_2_–** | 3,4-dichlorobenzyl | benzyl | 0 | 4 |
| 31 | **–SO_2_–** | 2,4-dichlorobenzyl | - | 1 | 47 |
| 32 | **–SO_2_–** | 3-fluorobenzyl | benzyl | 0 | 33 |
| 33 | **–SO_2_–** | 3-methylbenzyl | *sec*-butyl | 0 | 47 |
| 34 | **–SO_2_–** | 2,4-dichlorobenzyl | benzyl | 0 | 56 |
| 35 | **–SO_2_–** | 2-chlorobenzyl | benzyl | 0 | 8 |
| 36 | **–SO_2_–** | 3,4-dichlorobenzyl | *sec-*butyl | 0 | 49 |
| 37 | **–SO_2_–** | 3-methylbenzyl | - | 1 | 37 |
| 38 | **–SO_2_–** | 2,5-dimethylbenzyl | *sec*-butyl | 0 | 17 |
| 39 | **–SO_2_–** | 4-methylbenzyl | methyl | 0 | 6 |
| 40 | **–SO_2_–** | 2-fluorobenzyl | Methyl | 0 | 58 |
| 41 | **–SO_2_–** | Benzyl | *sec*-butyl | 0 | 30 |
| 42 | **–SO_2_–** | 4-chlorobenzyl | isobutyl | 0 | 5 |
| 43 | **–SO_2_–** | 2,6-dichlorobenzyl | *sec*-butyl | 0 | 18 |
| 44 | **–SO_2_–** | 2-methylbenzyl | isobutyl | 0 | 13 |
| 45 | **–SO_2_–** | 2-methylbenzyl | isopropyl | 0 | 49 |
| 46 | **–SO_2_–** | Benzyl | Methyl | 0 | 51 |
| 47 | **–SO_2_–** | 4-methylbenzyl | *sec*-butyl | 0 | 19 |
| 48 | **–SO_2_–** | 2-chlorobenzyl | *sec*-butyl | 0 | 53 |
| 49 | **–SO_2_–** | 4-methylbenzyl | isobutyl | 0 | 5 |
| 50 | **–SO_2_–** | 2,4-dichlorobenzyl | Methyl | 0 | 6 |
| 51 | **–SO_2_–** | 4-methylbenzyl | Benzyl | 0 | 15 |
| 52 | **–SO_2_–** | 3,4-dichlorobenzyl | Methyl | 0 | 10 |
| 53 | **–SO_2_–** | 2-chlorobenzyl | - | 4 | 61 |
| 54 | **–SO_2_–** | 4-fluorobenzyl | isobutyl | 0 | 20 |
| 55 | **–SO_2_–** | 3-chlorobenzyl | - | 4 | 61 |
| 56 | **–SO_2_–** | 3-chlorobenzyl | isopropyl | 0 | 51 |
| 57 | **–SO_2_–** | 3-chlorobenzyl | benzyl | 0 | 58 |
| 58 | **–SO_2_–** | 4-methylbenzyl | - | 1 | 62 |
| 59 | **–SO_2_–** | 2,4-dichloro | isobutyl | 0 | 54 |
| 60 | **–SO_2_–** | 2-methylbenzyl | *sec*-butyl | 0 | 45 |
| 61 | **–SO_2_–** | 3-chlorobenzyl | Methyl | 0 | 61 |
| 62 | **–SO_2_–** | 3-chlorobenzyl | *sec*-butyl | 0 | 43 |
| 63 | **–SO_2_–** | 2,5-dimethylbenzyl | isobutyl | 0 | 5 |
| 64 | **–SO_2_–** | 3-methylbenzyl | isopropyl | 0 | 44 |
| 65 | **–SO_2_–** | 3,4-dichlorobenzyl | isopropyl | 0 | 5 |
| 66 | **–SO_2_–** | Benzyl | Benzyl | 0 | 20 |
| 67 | **–SO_2_–** | 3-fluorobenzyl | Methyl | 0 | 56 |
| 68 | **–SO_2_–** | 2-fluoro-6-chlorobenzyl | isopropyl | 0 | 33 |
| 69 | **–SO_2_–** | Benzyl | - | 4 | 55 |

**Scheme S1. Synthesis of *(S)*-O30 (5').**

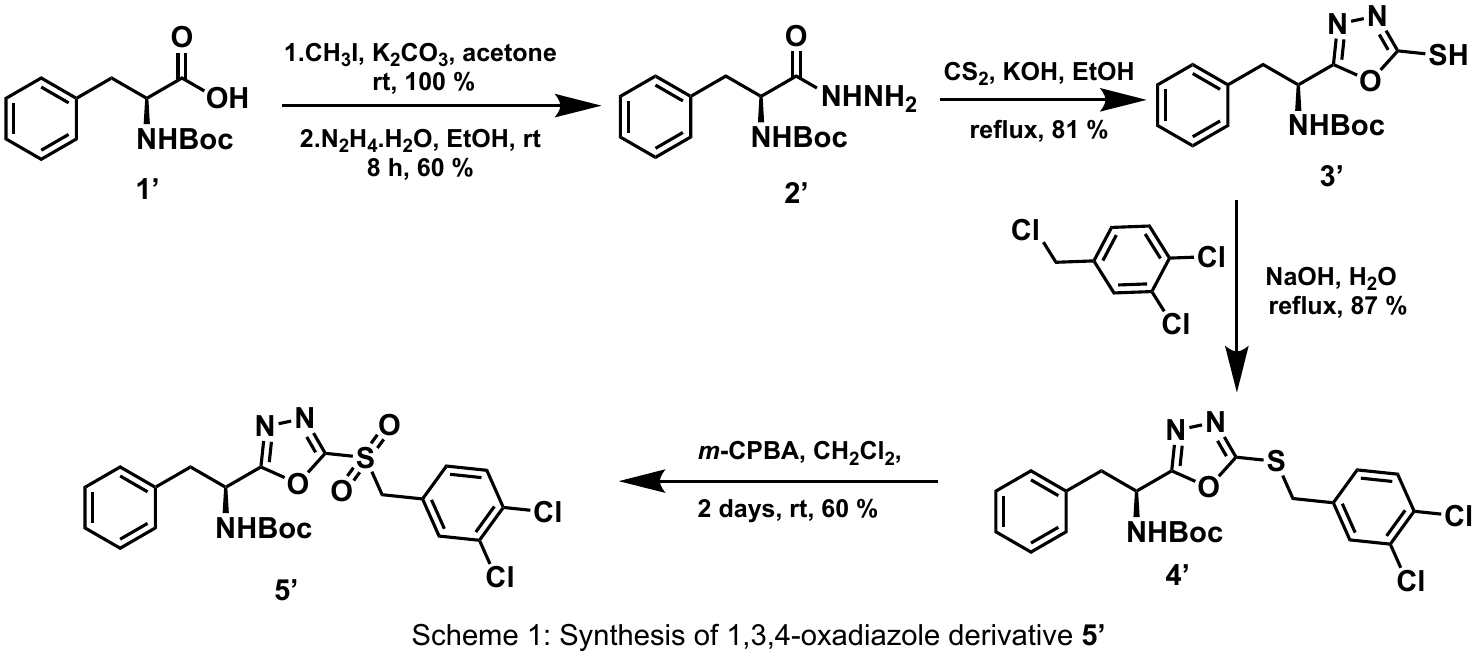

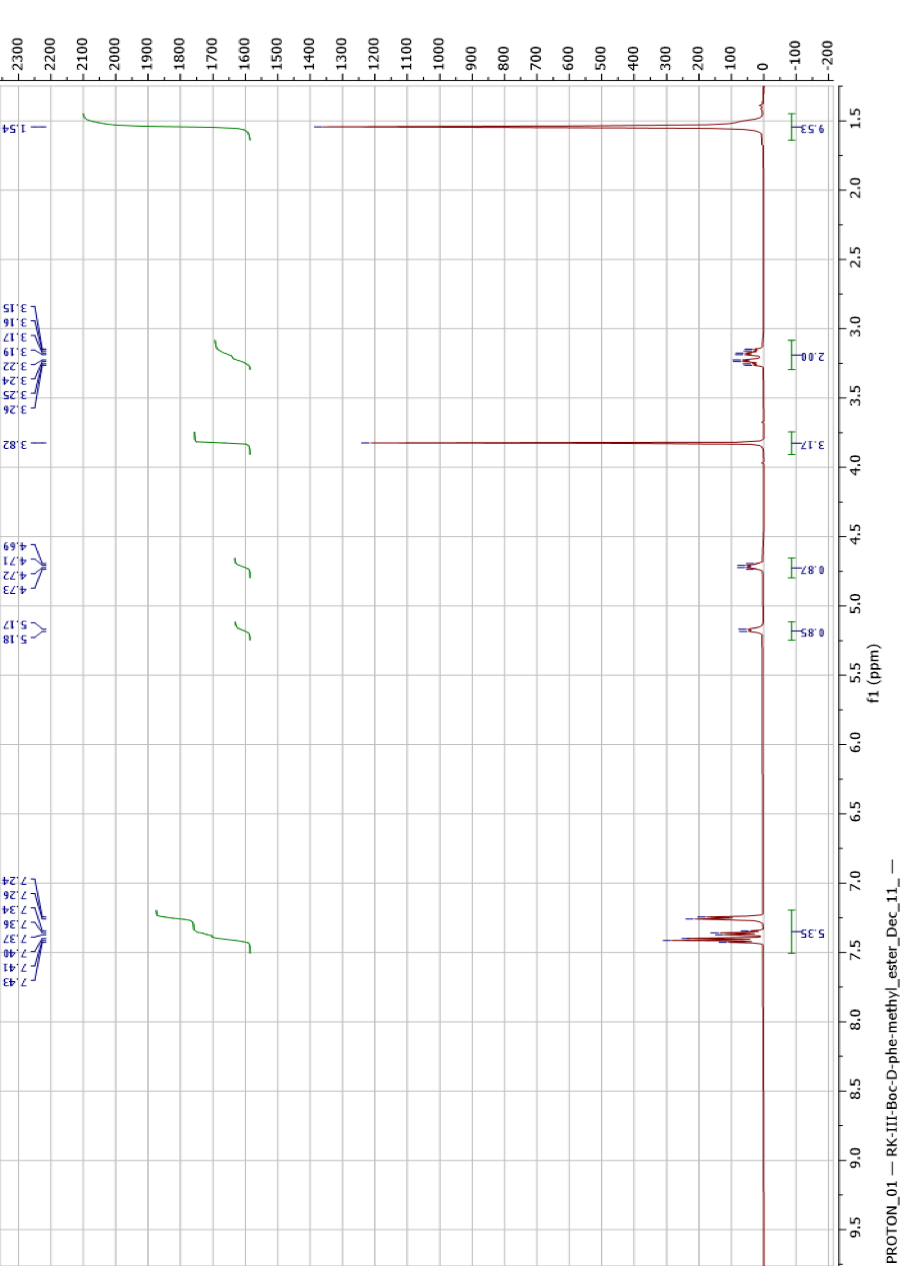
 **Figure S1.** ^1^H Spectrum of **1a** (500 MHz, CDCl_3_)

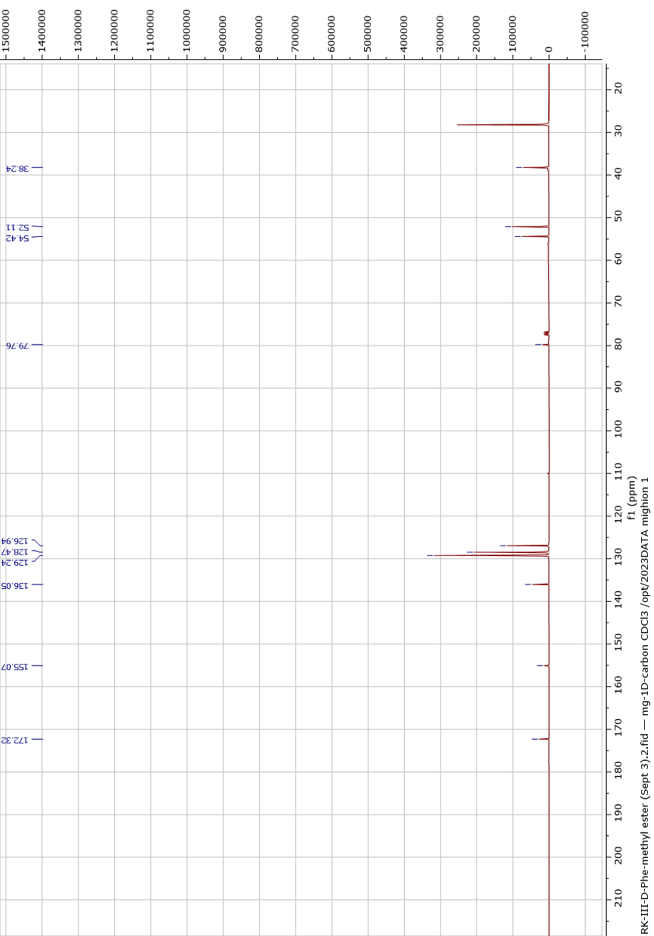
 **Figure S2.** ^13^C Spectrum of **1a** (101 MHz, CDCl_3_)

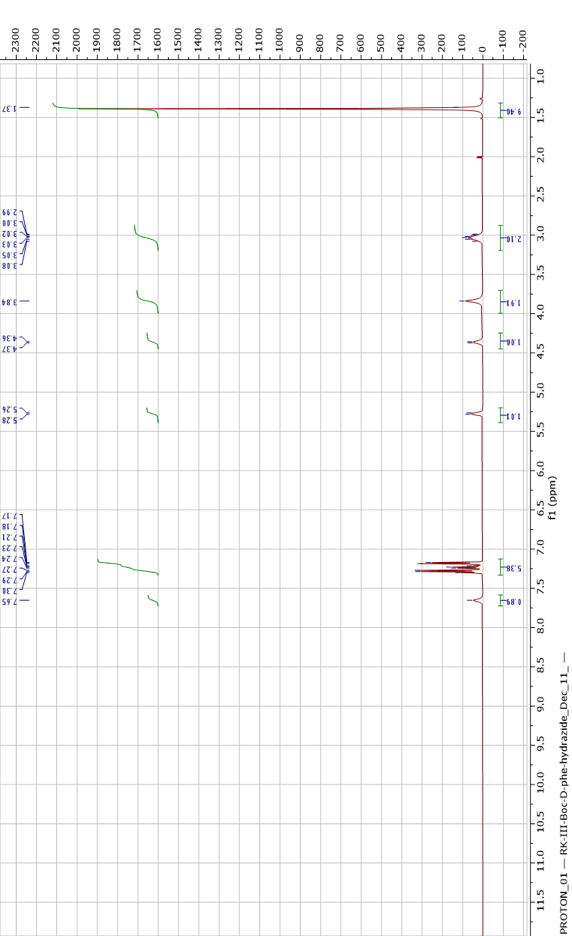
 **Figure S3.** ^1^H Spectrum of **2** (500 MHz, CDCl_3_)

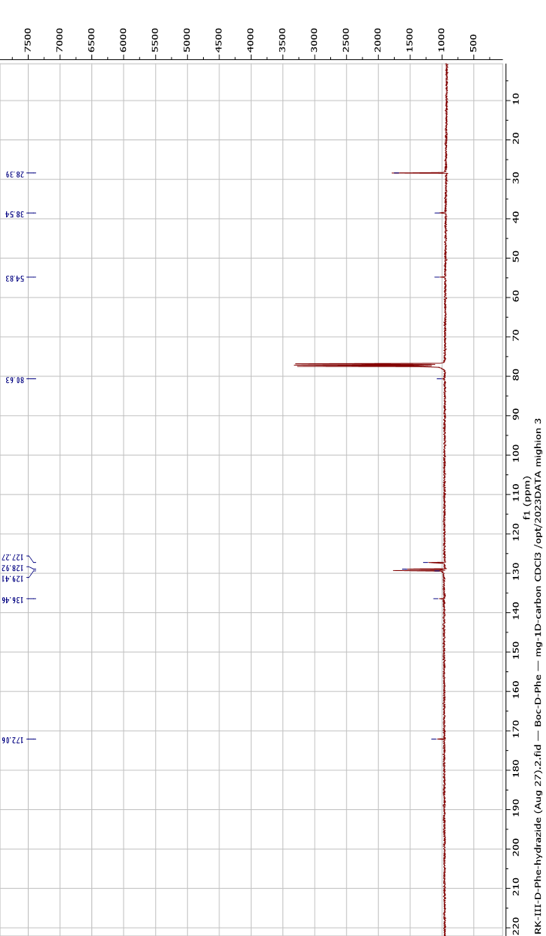
 **Figure S4.** ^13^C Spectrum of **2** (101 MHz, CDCl_3_)

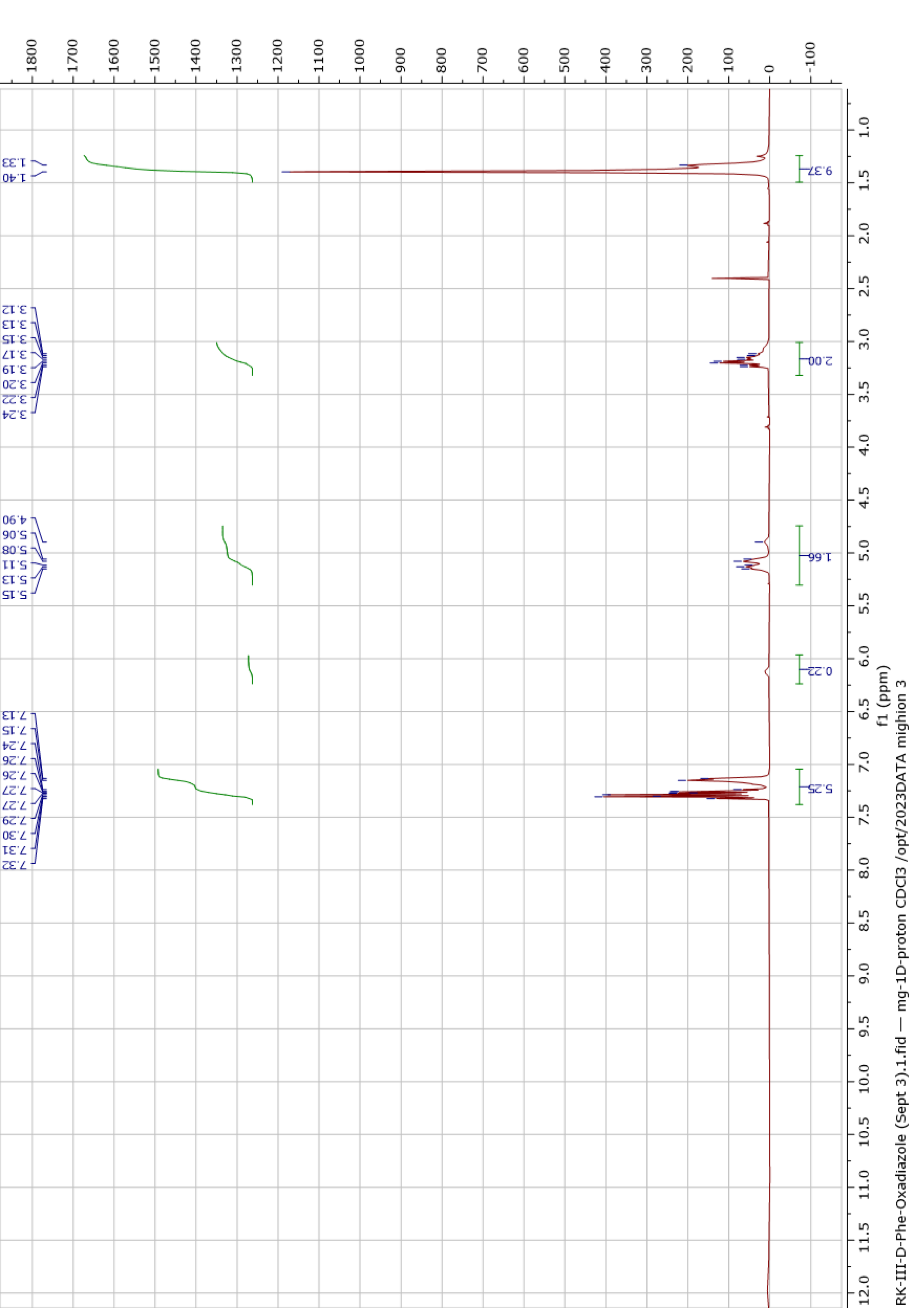
 **Figure S5.** ^1^H Spectrum of **3** (400 MHz, CDCl_3_)

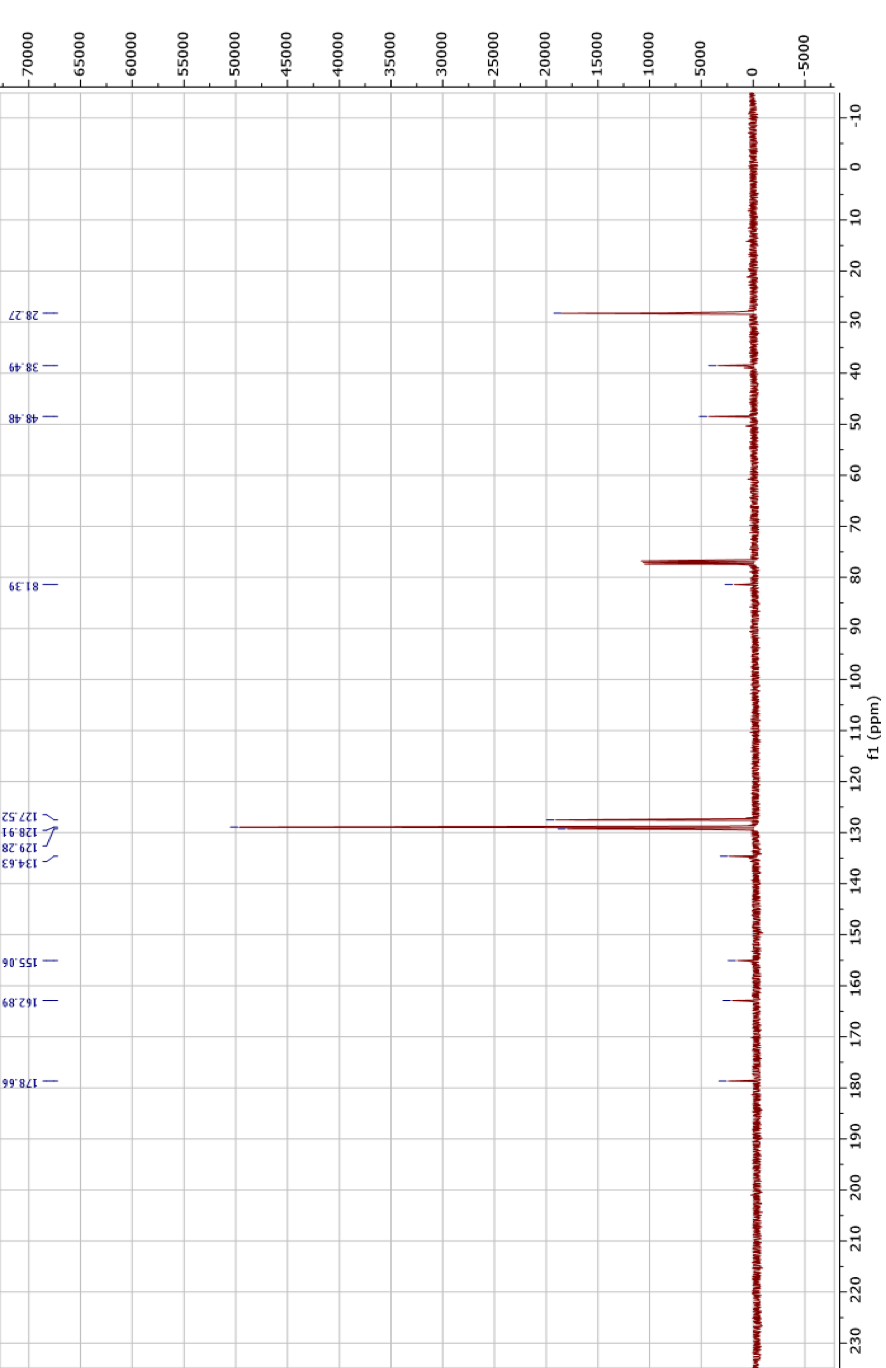
 **Figure S6.** ^13^C Spectrum of **3** (101 MHz, CDCl_3_)

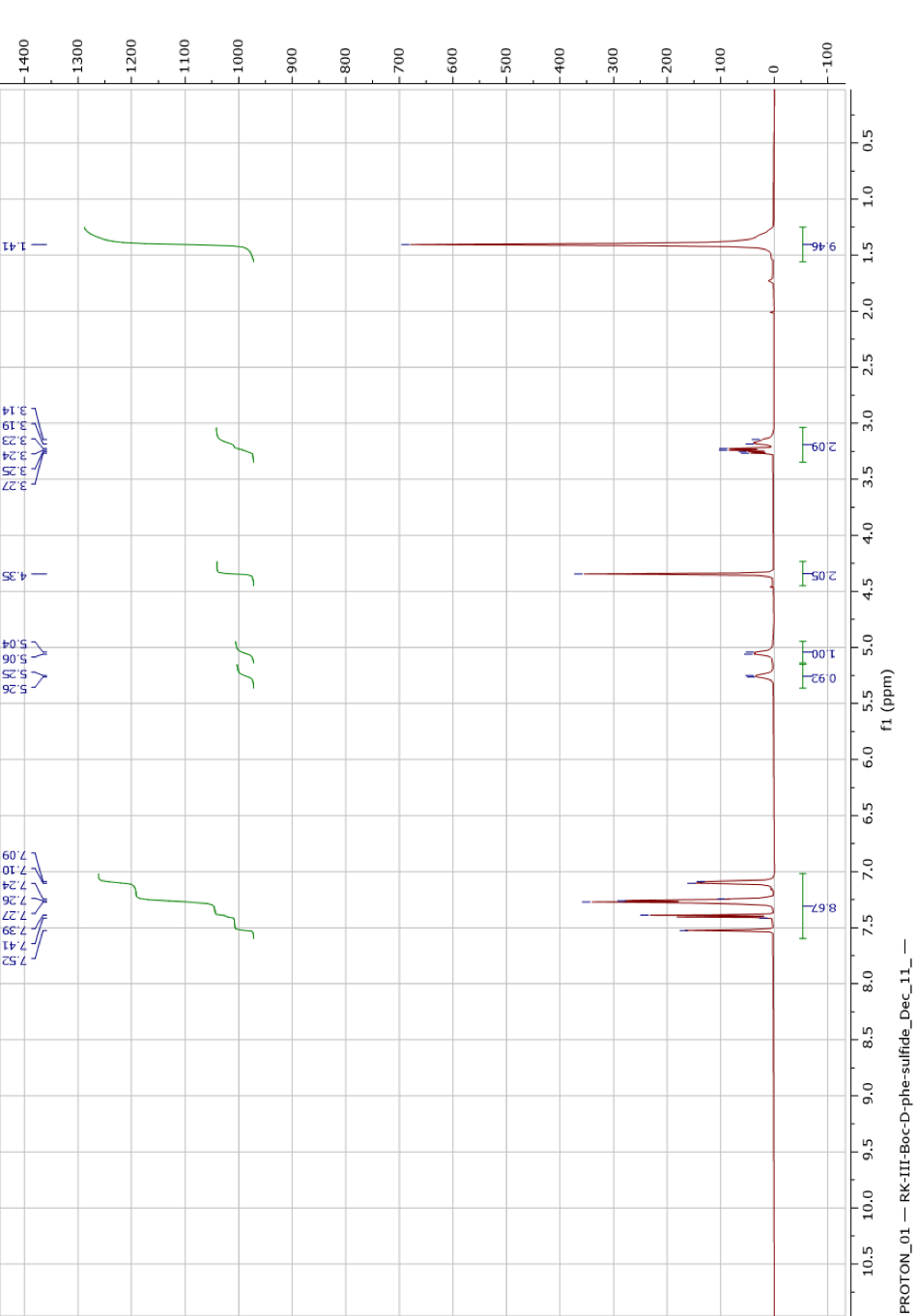
 **Figure S7.** ^1^H Spectrum of **4** (500 MHz, CDCl_3_)

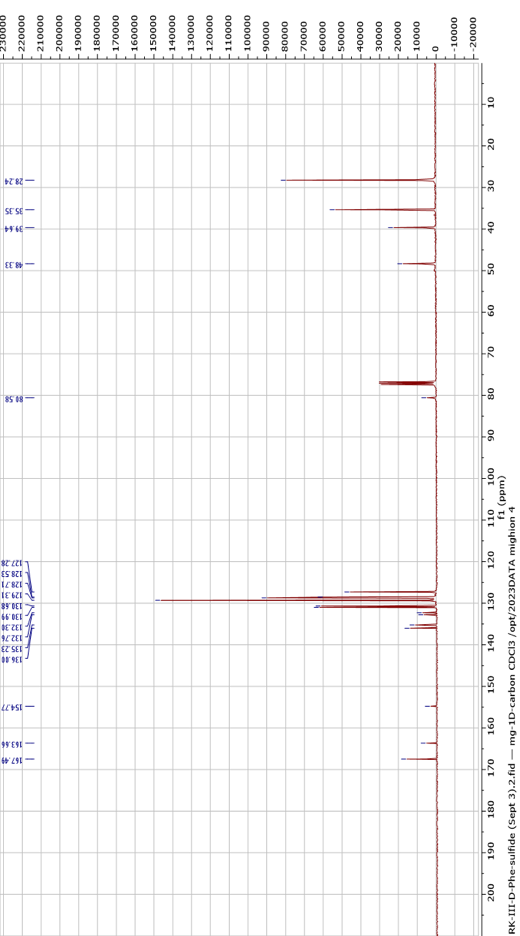
 **Figure S8.** ^13^C Spectrum of **4** (101 MHz, CDCl_3_)

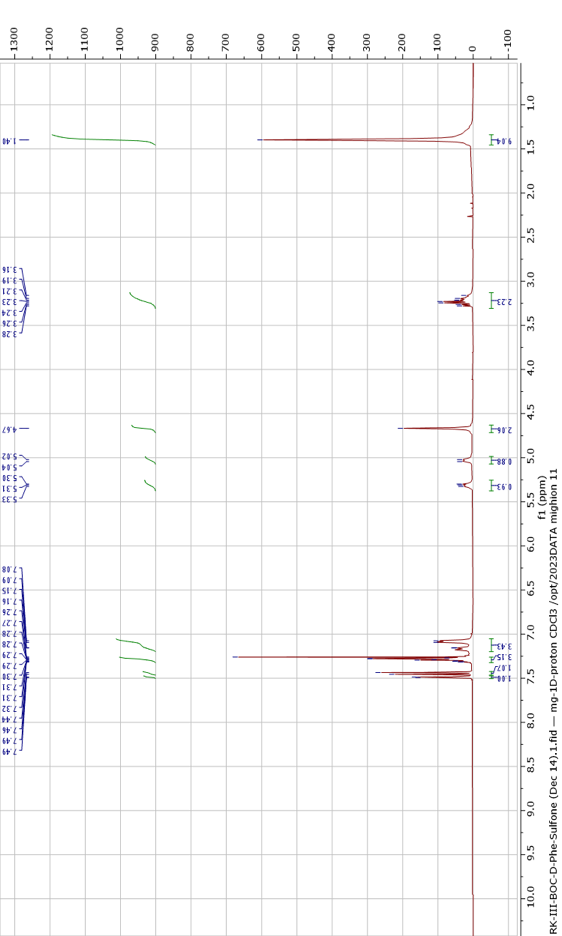
 **Figure S9.** ^1^H Spectrum of **5** (400 MHz, CDCl_3_)

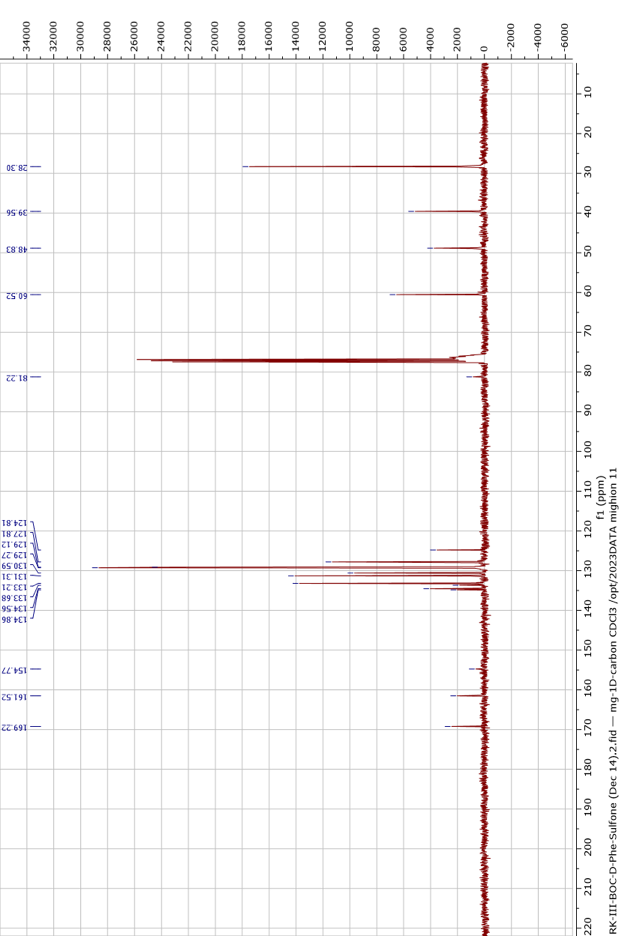
 **Figure S10.** ^13^C Spectrum of **5** (101 MHz, CDCl_3_)

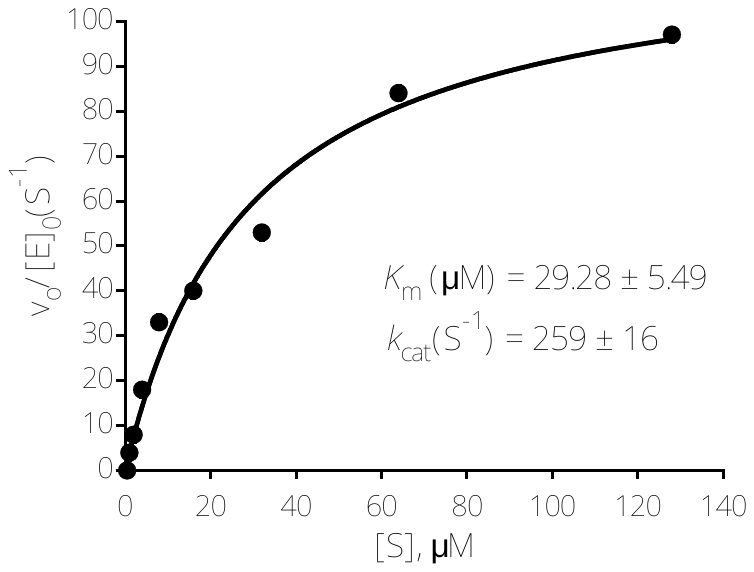

**Figure S11.**

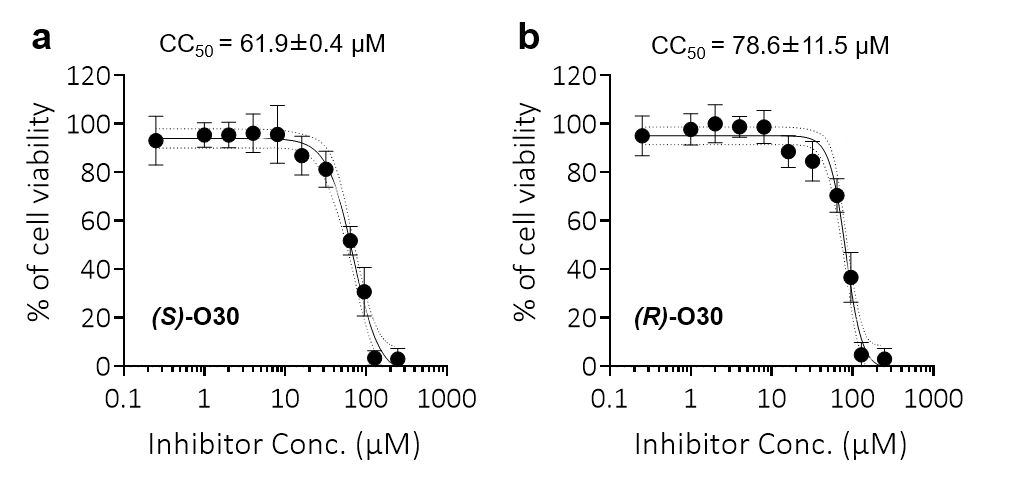

**Figure S12. Half-maximal cytotoxic concentrations (CC_50_) of both enantiomers against human HEK239 cells.**
